# From skylight input to behavioural output: a computational model of the insect polarised light compass

**DOI:** 10.1101/504597

**Authors:** Evripidis Gkanias, Benjamin Risse, Michael Mangan, Barbara Webb

## Abstract

Many insects navigate by integrating the distances and directions travelled on an outward path, allowing direct return to the starting point. Fundamental to the reliability of this process is the use of a neural compass based on external celestial cues. Here we examine how such compass information could be reliably computed by the insect brain, given realistic constraints on the sky polarisation pattern and the insect eye sensor array. By processing the degree of polarisation in different directions for different parts of the sky, our model can directly estimate the solar azimuth and also infer the confidence of the estimate. We introduce a method to correct for tilting of the sensor array, as might be caused by travel over uneven terrain. We also show that the confidence can be used to approximate the change in sun position over time, allowing the compass to remain fixed with respect to ‘true north’ during long excursions. We demonstrate that the compass is robust to disturbances and can be effectively used as input to an existing neural model of insect path integration. We discuss the plausibility of our model to be mapped to known neural circuits, and to be implemented for robot navigation.

**Author summary:** We propose a new hypothesis for how insects process polarised skylight to extract global orientation information that can be used for accurate path integration. Our model solves the problem of solar-antisolar meridian ambiguity by using a biologically constrained sensor array, and includes methods to deal with tilt and time, providing a complete insect celestial compass output. We analyse the performance of the model using a realistic sky simulation and various forms of disturbances, and compare the results to both engineering approaches and biological data.

## Introduction

Orientation cues are required for spatial behaviours from following a straight line [1] to migrating across continents [2] (for a theoretical proof see [3]). Idiothetic cues such as those generated in the mammalian vestibular system are useful for short time periods but are inherently problematic because of accumulating errors. To avoid this limitation, many animals, in particular insects, have developed an array of sensory systems to detect allothetic directional cues in their environments: magnetic (butterflies [4], moths [5], ants [6]), wind (moths [7], ants [8]), and visual [solar compass -(honeybees [9], crickets [10], locusts [11], butterflies [12], ants [13], dung beetles [14]); star compass - (dung beetle [15])]. The benefit of such accurate external compass systems is exemplified in the behaviour of desert ants, who utilise the sky polarisation pattern [16]. In a habitat with few if any landmarks, these ants can integrate the distance and directions travelled on a tortuous search path of up to a kilometre in length and make a direct return home when food is found [17]. Our main aim in this paper is to study the potential accuracy with which an insect can estimate its allocentric direction from the sky polarisation pattern, given realistic constraints on the environmental cues, the sensory system, and the various sources of disturbances.

The primary directional cue used for path integration by central place foraging insects such as desert ants [18] is the sky (sometimes called the celestial compass). The position of the sun (or moon) in the sky - as well as providing a direct directional reference point - determines the properties of light across the skydome including intensity and chromatic gradients, and a specific pattern of polarisation. Linear polarisation of light is the alignment of orientation of oscillation of the electromagnetic wave to a single plane. As light from the sun passes through earth’s atmosphere it undergoes a scattering process [19–22] producing differing levels of polarisation across the sky-dome relative to the position of the sun. From the point of view of an earth based observer, as the angular distance from the sun increases from 0° to 90°, the degree of linear polarisation in skylight increases, with the principal axis of polarisation perpendicular to the observer-sun axis, forming concentric rings around the sun. Angular distances above 90° have decreasing degree of linear polarisation [23].

The insect celestial compass has been studied extensively in the honeybee *Apis mellifera* [9, 24], the cricket *Gryllus campestris* [10, 25–27], the locust *Schistocerca gregaria* [11, 28], the monarch butterfly *Danaus plexippus* [12], the dung beetle *Scarabaeus lamarcki* [14] and the desert ant *Cataglyphis bicolor* [13]. Insects perceive polarised light through a specially adapted region of their upper eye known as the *dorsal rim area* (DRA). For the ommatidia in this area, the light sensing elements (microvilli) do not twist relative to each other, resulting in units that are sensitive to specific polarisation angles. Ommatidia in the DRA are connected to *polarisation sensitive (POL) neurons* in the *medulla* of the insect optic lobe which follow a sinusoidal activation profile under a rotating linear polarisation input [10]. The maximum and minimum activation is separated by 90^*°*^, consistent with an antagonistic input from at least two polarisation-sensitive channels with orthogonal *e-vector* tuning orientation (the e-vector is the electric vector component of the light’s electromagnetic energy and is orthogonal to the direction of propagation). The identity and spike-rate of each POL neuron thus encodes information about the *angle* and *degree* of polarisation respectively for the specific region of sky from which the associated ommatidia samples. An array of such sensing elements arranged appropriately may hence be sufficient to decode the sun position, without using any additional sky cues [29].

From the optic lobe the pathway for polarised light processing has been traced through several neuropils in the insect brain. Key processing stages for polarised light are the dorsal rim lamina and medulla, specific layers of the lobula, the anterior optic tubercle, and the lateral complex with giant synapses; before reaching a highly structured midline neuropil known as the central complex (CX) [28, 30]. A variety of neuron types within the CX region have been shown to have polarisation dependent responses, including CX inputs in the form of three types of tangential neurons [28] (TL1, TL2 and TL3) which synapse with columnar neurons [11, 31, 32] in the ellipsoid body/lower division of the central body. Most strikingly, intracellular recordings from neurons in the CX protocerebral bridge have revealed an orderly polarisation tuning preference across the eight columns, described as an internal compass [33]. More recently, the same structure has been observed through neurogenetic imaging in Drosophila to act as a ring attractor encoding the heading direction of the insect relative to a prominent visual cue [34]. However, there remains an inconsistency between these observations, as the polarisation tuning appears to range from 0°to 180° with each successive column tuned ∼22.5° degrees apart [33], whereas the fly’s compass covers at least 270°, changing by around ∼35° per column [34] and might be assumed to form a 360° representation of space [32].

In a recent model, we used anatomical constraints for processing within the CX to explain how compass information in the protocerebral bridge could be combined with speed information to carry out path integration, and subsequently steer home [35]. This model assumed a 360° compass across the 8 tangential cells (TB1) of the protocerebral bridge (PB) could be derived from sky polarisation cues. In this paper, we first determine whether in principle such a signal can be recovered from a simulated array of POL-neurons stimulated by a realistic sky polarisation pattern. We further investigate whether this signal can deal with, or be plausibly corrected for, potential disturbances such as partially obscured sky, tilting of the sensor array, and the movement of the sun with passing time. We evaluate the potential accuracy of compass information both in absolute terms, and in the context of path integration. Finally we show how the discrepancy in biological data for the protocerebral bridge tuning pattern [33, 34] might be resolved, by testing our model with artificial polarisation patterns.

## Materials and methods

### Overview

This study investigates how navigating insects can transform solar light into an earth-based compass signal that is sufficiently stable and accurate to drive precise path integration behaviour. Fig 1 provides an overview of the modelling pipeline. We start with a physical model of skylight, which is used as input to a biomimetic sensor array based on the desert ant eye. We then take a more direct computational approach to generate compass output from the insect eye input, by defining a hypothetical neural architecture that will reconstruct the sun position from this input, with additional mechanisms to correct for tilt and for passing time. As the precise neural connectivity underlying these transformations in the insect is unknown, this is a proof of principle that can provide hypotheses for future investigation of this circuit (see discussion). We then use the output of the compass as input to an existing biologically constrained model of path integration in the central complex, and test it in a closed loop agent simulation. The properties of our model are drawn from a variety of insects that are shown to have a celestial compass, as detailed in Table 1, but with a specific focus towards the desert ant.

**Table 1.** The cross-insect properties of our model.

| DRA layout |  |  | photo-receptors |  | optic lobe<br>neurons |
| --- | --- | --- | --- | --- | --- |
| $n$ | $\omega$ | shape | $\lambda_{\max}$ | $\rho$ | |
| 60 | 56° | fan-like | 350 nm | 5.4° | POL-OP |
| (desert ant [36, 37]) |  |  | (desert ant [38]) |  | (desert ant & cricket [39]) |

| tilt compensation | time compensation | compass in CX |
| --- | --- | --- |
| ocelli, visual horizon<br>(desert ant [40] & various insects [41]) | ephemeris function<br>(desert ant [42] & honeybee [43]) | heading code<br>(desert locust [33]) |

$n$ , number of ommatidia on the sensor; $\omega$ , receptive field of the sensor; $\lambda_{\max}$ , maximum spectral sensitivity (wavelength) of a single photo-receptor; $\rho$ , acceptance angle of a single photo-receptor. Specific species of insects for each property: **DRA layout**, *C. bicolor* & *C. fortis* [36], *C. albicans*, *C. bicolor* & *C. fortis* [37]; **photo-receptors**, *C. bicolor* [38]; **optic lobe neurons**, *C. bicolor* & *G. campestris* [39]; **tilt compensation**, *C. bicolor* [40], various insects [41]; **time compensation**, *C. bicolor* & *C. fortis* [42], *A. mellifera ligustica* L. [43]; **compass in CX**, *S. gregaria* [33].

**Fig 1.**
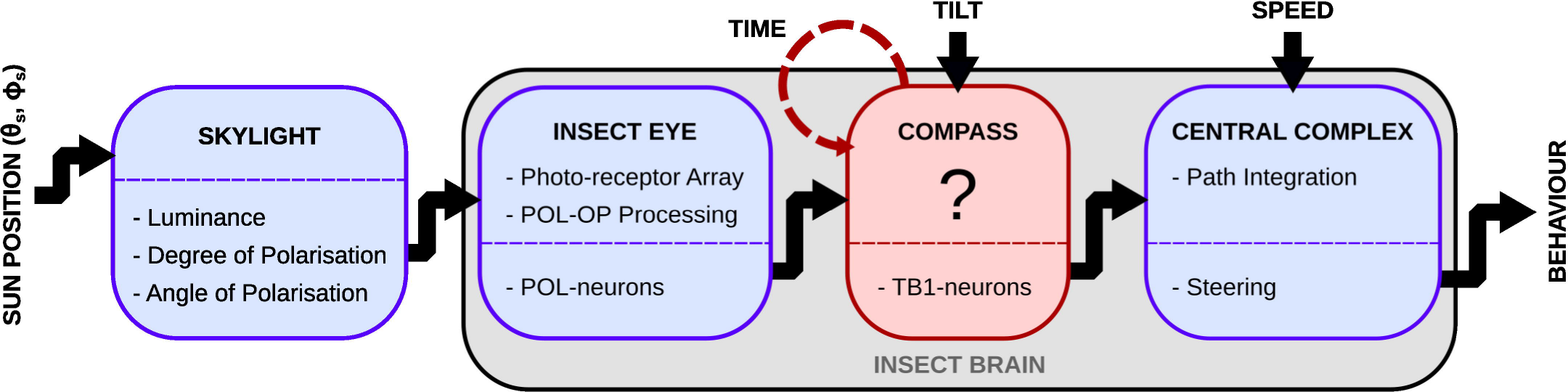
Overview of the modelling pipeline. Our simulation consists of four consecutive models (left to right). Given the position of the sun, a realistic skylight model [44] provides the predicted luminance, degree and angle of linear polarisation for every point in the sky-dome. This provides input to the eye model, based on biological data from insects [10, 38], which defines an array of polarisation sensitive photo-receptors facing different parts of the sky, and uses opponent processing to produce luminance-independent POL-neuron responses. The compass model provides a hypothesis for the unknown neural process that converts the POL-neuron response to a true compass signal in the TB1 neurons; this also utilises information about tilt of the sensor array, and allows for the movement of the sun with passing time. Finally, the compass neurons’ output is used along with speed as input to an anatomically grounded model of the central complex [35] which performs path integration and produces an output signal that can steer the insect back home. Blue boxes represent known systems in the pathway of the skylight; the red box represents a fuzzy/unknown system, which is the main focus of modelling in this paper.

### Skylight

To test our neural model, we need to simulate the incoming light using a skylight dome model. Previous computational studies of the insect POL-system have often copied the typical stimulus input from experimental studies, i.e., a rotating linear polariser [45]. However, the topology of the ommatidia and the neural processing of the compass system in the insect brain have evolved under real sky conditions, hence using a more realistic input can be critical to understanding the function. Specifically, as we will show, the real sky pattern breaks the symmetry conditions that inherently prevent 360° directions being recovered from a simple linear polarisation cue.

We use the skylight dome model described in [44], which gives a very realistic luminance and linear polarisation information pattern (a sample of its output is illustrated in Fig 2). This model is the most accurate description of the skylight distribution currently available, and for a detailed description we refer the reader to the original work [44] which we follow directly; a brief description is also given in the section S3 Appendix outlining the objective function used in our evaluation. Given the position of the sun and a set of points in the sky, this model generates the *luminance*, *degree* (DOP) and *angle of polarisation* (AOP) for those points. Tuning by geo-referenced input parameters allows realistic sky patterns to be estimated for specific locations. Therefore, plugging into the model the location from our own desert ant fieldwork site (Seville, Spain) allows us to run simulated experiments for desert ants. This way, we can study the response of their POL-sensitive neurons using near natural stimuli.

**Fig 2.**
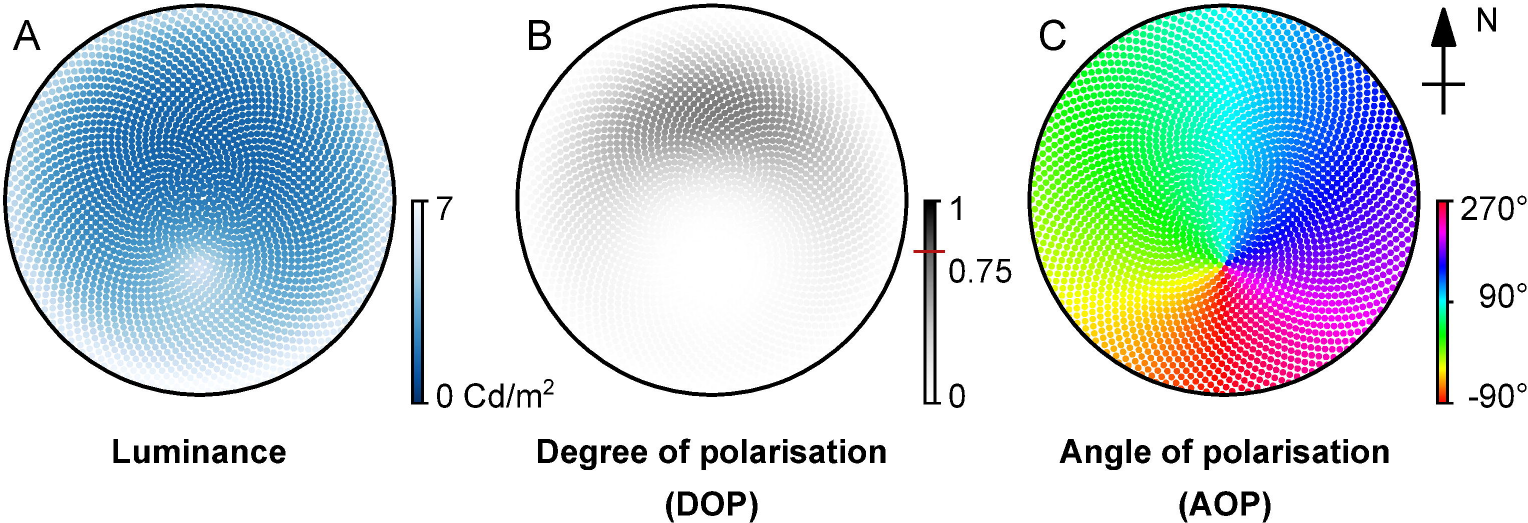
Sample output from the skylight dome model [44]. The luminance pattern of the sky is proportional to its intensity and describes the amount of light per area unit existing in a specific direction. Along with the chromaticity coordinates, it can provide spectral information. (B) The degree of linear polarisation pattern in the sky based on the scattering of the light on atmospheric particles. It is defined by the fraction of the polarised portion over the total intensity. The red line on the colour-bar showing the *d* = 0.75 indicates the maximum DOP observed in the skylight simulation. (C) The angle of polarisation pattern in the sky is defined by the average e-vector (electric part of an electromagnetic wave) orientation of the photons. The black circle in the figures denotes the horizon. In all panels the sun position is 30° south.

### The Insect Eye

#### Dorsal rim ommatidia

We developed a simulated sensory unit to mimic the function of the ommatidia in the DRA of the desert ant (Fig 3A and B). We consider each ommatidium to be an individual sensorial unit that can be freely arranged in space to match the compound structure of the real ant’s DRA (Fig 3D and E).

**Fig 3.**
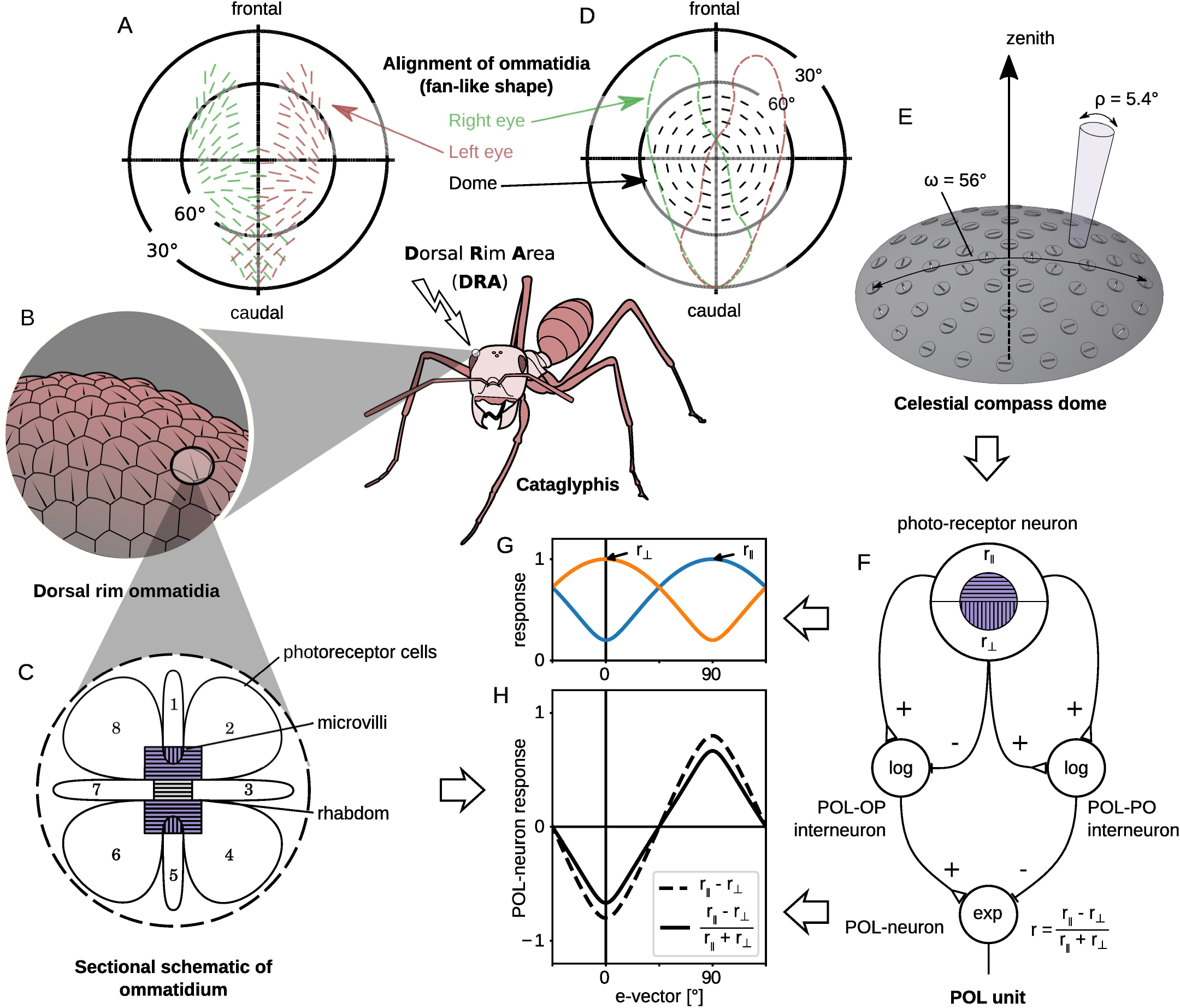
Processing stages of light in the biological and artificial DRA. (A) Top view of the fan-like arrangement of the ommatidia on the Cataglyphis DRA for both the right (green) and left (red) eyes; adapted and modified after [36]. (B) A closer look at the DRA, which is composed of hexagonal ommatidia. (C) An ommatidium on the DRA of the compound eye of the *Cataglyphis* has 8 photo-receptor cells, with parallel microvilli direction in 2, 3, 4, 6, 7 and 8, and perpendicular in 1 and 5; the colour violet indicates sensitivity to ultraviolet light. (D) Top view of the fan-like arrangement of the units on our sensor. The dashed lines show the overlap with the areas of the left (red) and right (green) Cataglyphis DRAs. (E) 3D representation of the sensor array in the eye model with visual field *ω* = 56°: the 60 discs on the dome are different units (ommatidia) with acceptance angle *ρ* = 5.4°; the orientation of the lines on the circles denote the direction of the main (parallel) polarisation filter. (F) Model of a POL-unit: the photo-receptor neurons combine a UV-sensor (photo-receptor) and a polarisation filter (microvilli), and have a square-root activation function. The normalised difference of the photo-receptor neurons is calculated by the POL interneurons. The empty triangular and dashed synapses denote excitatory and inhibitory connections respectively. (G) Simulated response of the two photo-receptors in one unit in partially linearly polarised light of intensity *I* = 1 and degree of polarisation *d* = 0.9 against different e-vector orientations. (H) Simulated response of the POL-neuron to the input of Fig 3G; the dashed line shows the response of the POL-OP interneuron, and the solid line is the response of the POL-neuron (normalised difference). B and C figures adapted and modified from [46]. F, G and H are after [10].

For an insect, the acceptance angle of the ommatidia affects the volume of skylight perceived and the maximum polarisation contrast sensitivity. This is defined by the optical properties of the cornea and the crystalline cones. Light passing through these lenses is focused through a light-guide onto 8 light sensitive cells (rhabdoms) that have a preference for one of two groups of perpendicularly oriented microvilli, that work as linear polarisation filters (Fig 3 C).

For each simulated ommatidium in our model we define a fixed acceptance angle, *ρ* = 5.4°, and spectral sensitivity, *λ*_max_ = 350 nm wavelength, based on the ant eye [38] and use 2 perpendicularly arranged photo-receptor channels. We notate *s*_||_ (*s*_⊥_) the stimulus of the photo-receptor channels that have parallel (perpendicular) aligned polarisation filters with respect to the ommatidium orientation. These follow a sinusoidal response curve to the e-vector orientation with the highest value when fully aligned with the preferred polarisation angle and the lowest when perpendicular. The photo-receptor neuron transforms this raw input using a square-root activation function, 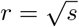 which reduces right skewness, transforming the sinusoid as shown in Fig 3G. This transformation is essential in order to penalise the high illuminations of the bright sky. It acts similarly to the logarithmic transformation *r* = log(*s*), introduced by [39] and used on the Sahabot robot [47]. The main advantage of the square-root transformation is that it can be applied to zero values (darkness or linearly polarised light perpendicular to the filter orientation). As the exact transformation of the photo-receptors signal in the insect’s eye remains unknown [48], any transformation that reduces the right skewness could be theoretically correct.

#### POL-neurons: Polarisation contrast

The first stage of the global-orientation encoding in insects is performed by the *polarisation* units in the medulla of the optic lobe [10]. These units encode polarisation information from different points of the insect’s environment, creating a polarisation map. We follow the approach described in [39] to define how light perceived from our sky model is transformed into POL-neuron activity. This produces luminance insensitivity in each POL-neuron, capturing only the polarisation contrast in specific e-vector directions.

The photo-receptor cells propagate their response to the POL-neuron output via two interneurons (see Fig 3F). The *polarisation-opponent* (POL-OP) interneuron (left) computes the difference between the responses of the photo-receptor neurons, *r*_OP_ = *r*_||_ − *r*_⊥_, similarly to the output (POL-neuron) in [47]. The *polarisation-pooling*(POL-PO) interneuron (right) approximates the overall intensity of the input light, by pooling the responses of the opponent components, *r*_PO_ = *r*_||_ + *r*_⊥_, and is used as a normalisation factor that removes this luminance information from the signal. Both interneurons propagate the logarithm of their output signal to the POL-neuron.

The POL-OP interneuron excites the output POL-neuron, while the POL-PO interneuron inhibits it. As a result the activity of the output neuron is independent of the luminance. This feature is very important, as the activity of the POL-neurons in insects’ optic lobe is not correlated to the light intensity [13, 36]. This has been previously modelled by normalising the response of POL-neurons using the values measured from different directions [25, 47, 49, 50] (scanning model). Finally, the output neuron transforms the response using an exponential activation function in order to bring back the sinusoidal shape:

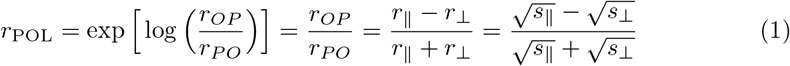

The above set of interneurons, along with the photo-receptors and the output POL-neuron compose a POL unit (Fig 3F). Fig 3G illustrates the response of the antagonistic photo-receptor neurons in a POL unit when the perceived light is partially and linearly polarised (degree of polarisation, *d* = 0.9, i.e. higher than found in skylight, but possible in experimental situations). The dashed line in Fig 3H illustrates the response of the POL-OP interneuron and the solid line shows the normalised POL-neuron response after dividing (inhibiting) by the POL-PO signal.

The output signal of our POL-neuron thus matches the one found in ants’ and crickets’ POL neurons [39]. However, we note there is no specific evidence for the hypothesised interneurons in this model.

#### Dorsal Rim Area: Layout

The layout of the sensor approximates a joint DRA (from both compound eyes; see Fig 3A) of an ant, leading to a cyclopic DRA. We approximate the sampling pattern of the cyclopic DRA by homogeneously distributing simulated ommatidia on the dome using the *icosahedron triangulation* method (Fig 3D), as widely used in computer graphics for sphere representations (detailed description of the design of the sensor in S1 Fig and a table with the spherical coordinates of all the POL units on the dome in S2 Table). The resulting pattern of polarisation preferences in our population of simulated ommatidia matches the overall specifications of the fan-like shape reported in ants [36–38] (Fig 3A). More specifically, as Fig 3E indicates, each ommatidium, with *acceptance angle* ρ = 5.4°, is aligned to its respective concentric notional ring centred at the zenith of a dome. Using the above method, we place *n* = 60 ommatidia on a dome-shaped surface, and the outer notional ring with radius 28° results in an *ω* = 56° receptive field for the whole eye model. Note that the view of each ommatidium can partially overlap with its neighbours. The *receptive field* is around half that of the ants’ DRA on the frontocaudal axis (see Fig 3D), and the total number of ommatidia are similarly 50% that of the real animal, keeping the resolution approximately the same.

The output of the ant eye model is a population (*n* = 60) of neurons closely matching the known characteristics of the POL neurons of the medulla of the crickets’ optic lobe [10]. This population thus forms the biologically constrained input layer for our visual processing model.

### The Compass

We built a *compass model* to transform the responses of the POL-neurons into the desired activation patterns of the TB1-neurons used for path integration. Alhough the anatomical pathway is known [51], the neurobiological processes on this pathway are as yet uncertain, so we have taken an information processing approach: given biologically realistic POL-neuron responses gathered from the skylight simulation we examine whether this input provides enough information for a biomimetic central complex model to drive steering. Fig 4 shows an overview of the model. The connection of POL-neurons to SOL-neurons in the *solar layer* implements a *sum-of-sinusoids* model to recover an estimate of the solar azimuth. The *gating function* adjusts the connection weights to compensate for tilting of the sensor array. The *true compass layer* uses the confidence of the estimate to predict and compensate for the changing sun position over time. We describe each of these steps in more detail below.

**Fig 4.**
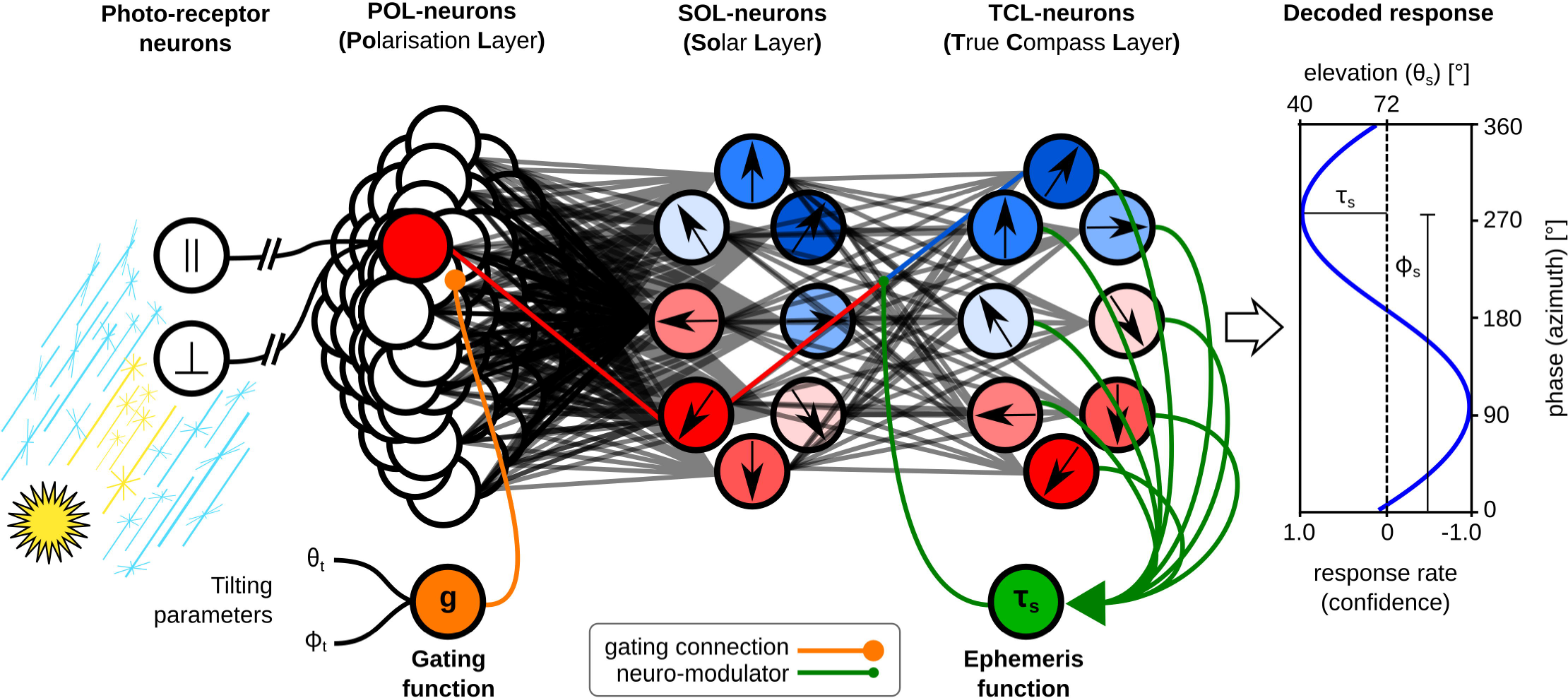
Overview of the compass model. The light information stimulates the photo-receptor neurons. Using the eye model, this is transformed to a POL-neuron population code. Tilting information (orange) is propagated through the gating function and creates a mask for the POL-neurons’ response, altering the relative weighting of information from different parts of the sensor array (see Fig 5). By combining information across the array, the response of SOL-neurons encodes an estimate of the solar azimuth and elevation [black arrows indicate the 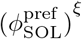 direction]. The TCL-layer uses the elevation information along with passing time as an ephemeris function (green) which modulates its input weights to rotate the SOL-neuron output and provide a true north estimate [black arrows indicate the 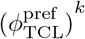 direction]. The response across the TCL-neurons is in the form of a sine-wave which can be decoded to determine an estimate of the azimuth (phase) and of the confidence of that estimate (amplitude). Note however that CX path integration circuit can use the TCL-neuron activity directly, without this explicit decoding.

#### Sum-of-sinusoids

The basic computation of the model is a *Sum-of-Sinusoids*, where the input is the 60 POL-neuron responses and the output is the position of the sun represented in an 8-neuron population code, named the *Solar Layer* (SOL). Each POL-neuron is connected to all SOL-neurons with a sinusoidal weighting function that represents the difference between the azimuthal direction of that POL-neuron’s receptor in the sensor array, 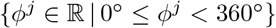, and the preferred direction of the SOL-neuron 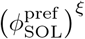, where 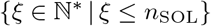 is the index of the *ξ*^th^ SOL-neuron. Thus each SOL-neuron sums a set of sinusoids with the same frequency, different phases (depending on *ϕ*^*j*^), and different amplitudes (depending on the POL-neurons activity), resulting in another sinusoid. Specifically, the response of the neurons of the solar layer is given by the equation below,

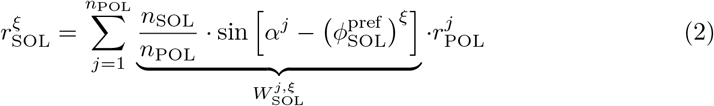

where *n*_POL_ = 60 and *n*_SOL_ = 8 are the number of POL- and SOL-neurons respectively,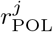 and 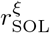 are their responses and 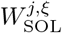 are the weights of the synapses connecting them. The weight depends on the orientation of the primary axis of the *j*^th^ ommatidum,*α*^*j*^ = *ϕ*^*j*^ − 90°, the preference angle of the respective *ξ*^th^ SOL-neuron, 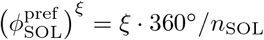, and the number of POL- and SOL-neurons. The more POL-neurons we have in the system the lower the synaptic weight connecting them to the SOL-neurons (smaller contribution), but the more SOL-neurons we have the higher the synaptic weight has to be (bigger contribution) amplifying the signal. Without this scaling factor the signal is very weak and sometimes vanishes.

Effectively, this equation activates each SOL neuron in proportion to the degree of polarisation opposite its preferred direction. As the degree of polarisation is maximum at the *cross-solar* point, the response of the SOL-neurons contains information about the predicted solar azimuth, and the confidence of this prediction. By ‘cross-solar’ we mean the point that is 90° away from the sun, in the solar-antisolar meridian, passing through the zenith point. For analysis, we can decode the population code using Fast Fourier Transform (FFT),

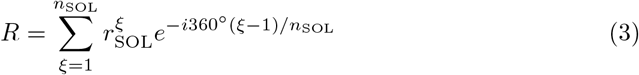

where 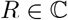. The angle of this complex number gives the estimated solar azimuth, 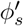, while the magnitude imply the confidence of this prediction, *τ*_*s*_,

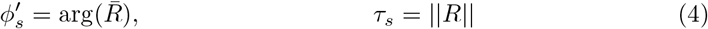

were arg(*⋅*) is the argument of a complex number, which gives the direction of the vector represented by it, and 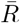 is the complex conjugate of *R*. The confidence, *τ*_*s*_, is just a factor and has no unit but it can be used to calculate the standard deviation, 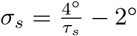, which is in degrees (see Results). Note that the insect does not need to extract the solar azimuth explicitly in this manner to be able to use the SOL encoding in further processing, but it may need to extract the confidence factor as discussed later (see Time compensation mechanism).

#### Tilt compensation mechanism

The above computation of azimuth assumes the cyclopic DRA is aligned with the sky dome, i.e. its zenith point is always pointing towards the sky zenith and it is laterally aligned to the horizon. In nature, the head of the animals may not remain aligned to the horizon, particularly in walking animals such as ants, which do not fully stabilise their head position [52]. As the head deviates from the horizon, the predictions of the above model become less and less accurate. To compensate for this error because of tilting, we added a gating function that receives information about the sensor tilt and modulates the response of the solar layer (see Fig 4 orange connections).

Specifically, the gating function uses the known head-tilt angle (see Discussion for where this may come from for the animal) to preferentially select inputs from ommatidia facing towards the most interesting region in the sky, in the form of a Gaussian ring shape centred on the zenith point (see Fig 5):

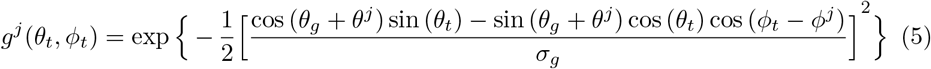

where *θ*^*j*^ and *ϕ*^*j*^ are the celestial coordinates (elevation and azimuth) of the relative position of the respective ommatidium, and (*θ*_*t*_, *ϕ*_*t*_) the celestial coordinates of the titling point. We notate *δ* = 90° − *θ*_*t*_ the tilting polar angle, which is the tilting angle with respect to the pole (zenith), such that zero tilt corresponds to the intuitively logical position of the sensor being centred on the zenith. The radius of this ring, *θ*_*g*_ = 40°, denotes the dominant focus direction of the system, i.e. where the peak of the Gaussian bump should be placed. The width of the ring (variance), *σ*_*g*_ = 13°, stands for the soft elevational receptive field for the SOL-neurons, i.e. the smoothing parameter. The above parameter values are the result of a global optimisation procedure (see Results).

**Fig 5.**
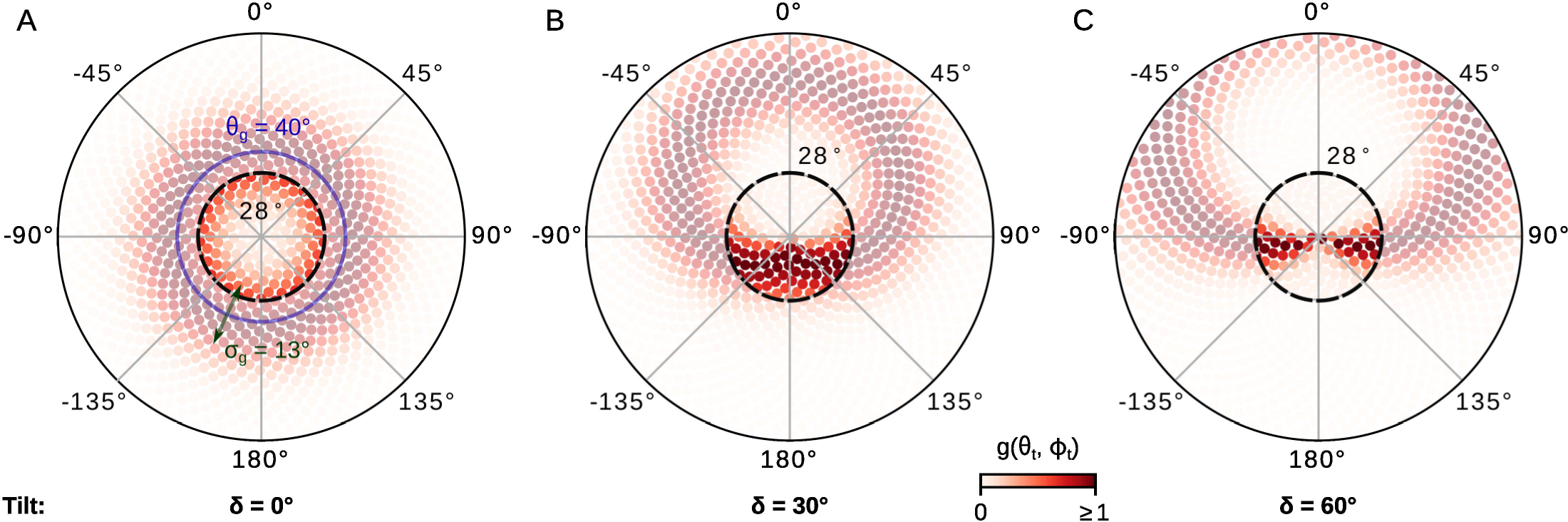
The gating function that compensates for tilt. The differing weightings of ommatidia input under three levels of tilt [(A) *δ* = 0° (*θ*_*t*_ = 90°), (B) *δ* = 30° (*θ*_*t*_ = 60°) and (C) *δ* = 60° (*θ*_*t*_ = 30°)] are shown, with darker shading indicating higher weighting. The inner dashed black circle delineates the actual receptive field of the simulated sensor (28° radius, equivalent to *ω* = 56° receptive field). The extended array (greyed out units) illustrates how this weighting adheres to a Gaussian function defined on the sky dome. The blue circle shows the dominant focus of the sensor (*θ*_*g*_ = 40°), and the green arrow shows the smoothing parameter (*σ*_*g*_ = 13°).

The sum-of-sinusoids from the previous section then changes to:

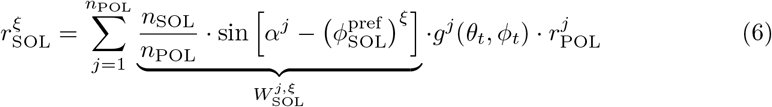

#### Time compensation mechanism

We extend the above model to provide a method by which insects might compensate for the changing celestial sky pattern over the course of the day and seasons. In particular, the aim is to provide a stable geocentric compass even though the solar azimuth varies compared to the true north. This is usually assumed to require an internal clock and calculation or learning of the ephemeris function [42, 43]. Here we suggest a possible solution that uses the current polarisation information only to estimate the ephemeris function and thus continuously adjust the compass. This is based on the observation that the confidence of the estimate of the solar azimuth is related to the solar elevation (see Discussion), from which we can infer the rate of change of the azimuth.

We add an extra layer to our model, the *true compass layer* (TCL), which is a copy of the SOL but the preference angle of its neurons changes through time (see Fig 4). The basic response of this layer is given by:

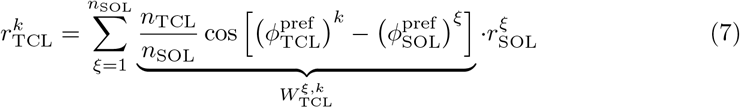

where 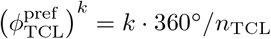 is the preference angle of the *k*^th^ TCL-neuron and *n*_TCL_ = 8 is the size of TCL-population.

As described earlier, we can derive from this neural representation the direction of the sun, 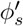, and the confidence of the prediction, *τ*_*s*_. In a low-disturbance environment and for a specific set of responses, the confidence can be transformed to a prediction of the solar elevation using the function below:

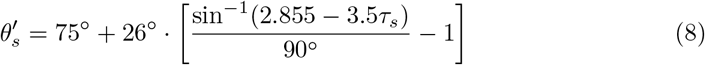

with domain 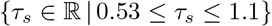 and range 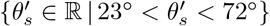 (see Fig 6A). For values of *τ*_*s*_ outside its domain, this equation would give the respective highest or lowest possible 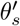. The above equation has been derived heuristically and it is a very simplified approximation of the inverse of the *equation of time* or *astronomical equation* [53]. When the confidence is high enough the predicted elevation, 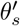, can be used to approximate the change rate, in degrees per hour (°/h), of the solar azimuth using the following equation (illustrated in Fig 6B):

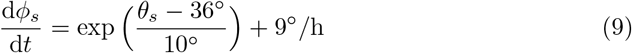

Integrating the above equation through time and using the information that the sun is always moving clockwise, we can approximate the shift of our predicted solar azimuth, 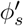, required to get a global orientation (see Fig 6C). This is implemented by introducing an update rule for the TCL-layer:

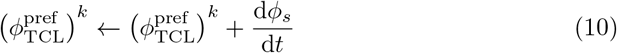

which updates the connections between the solar layer and the TCL-neurons through a recurrent connection (see Fig 4 green connections). Fig 7 shows the response and weights for all the different layers along with the corrections of the global direction when we add the tilt and time compensation mechanisms. The above integration gives an alternative answer to the question of how insects can develop their *ephemeris function* without using a clock.

**Fig 6.**
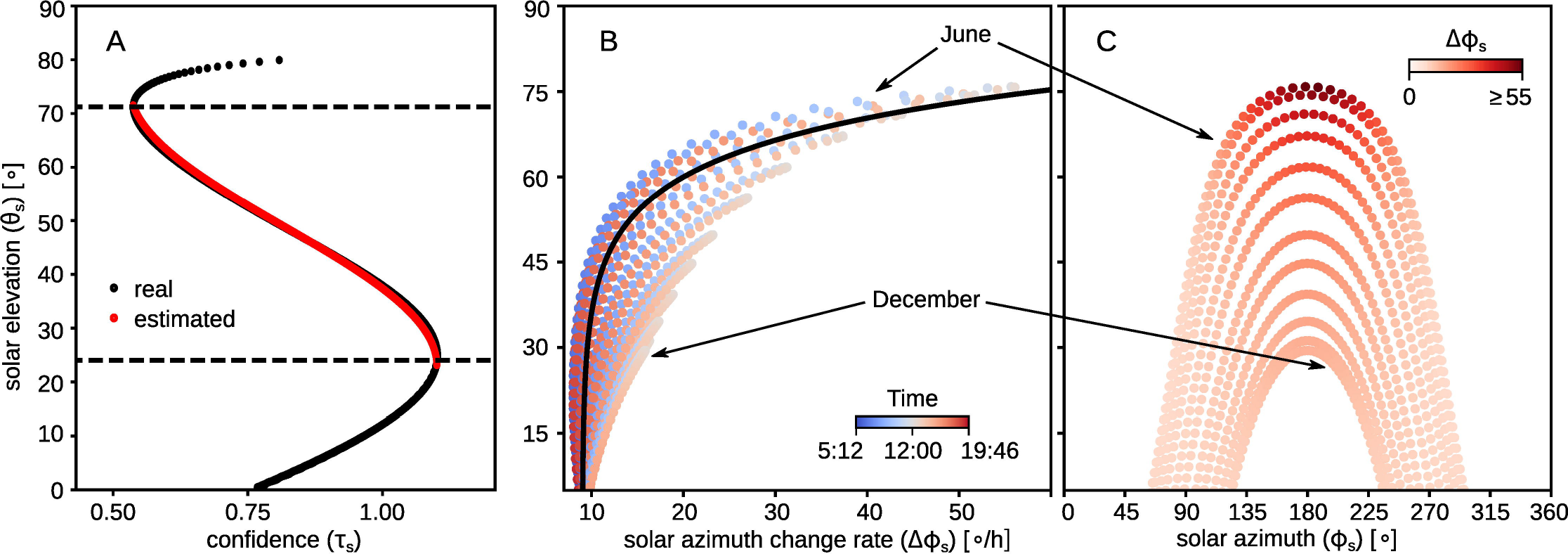
Using confidence of the estimate to compensate for time. (A) The confidence value of the compass response varies with the solar elevation (black dots). Within the range 23° − 72° this relationship can be used to estimate the elevation using Eq (8) (red dots). (B) The rate of change of the solar azimuth over time depends on the elevation [coloured dots represent different times of the day from morning (blue) to the evening (red)] and can be approximated using Eq (9) (black line). (C) Showing the solar elevation with respect to the solar azimuth for different times of the year. Each of the 12 imaginary curves in B and C correspond to the 21^st^ of every month; the sampling rate in each day is every 10 minutes from sunrise to sunset.

## Central Complex

The TCL-neuron output described above provides the required compass input that was assumed to be available in a previous anatomically constrained model of path integration in the insect central complex [35]. In that model, a set of 8 TB1 neurons with preferred directions {*k* • 45° | *k* = 1, …, 8*}* were activated with a sinusoidal relationship to the heading of the agent. We thus use an exact copy of this CX model, replacing its idealised input with our polarisation-derived compass signal to test its efficacy and robustness.

**Fig 7.**
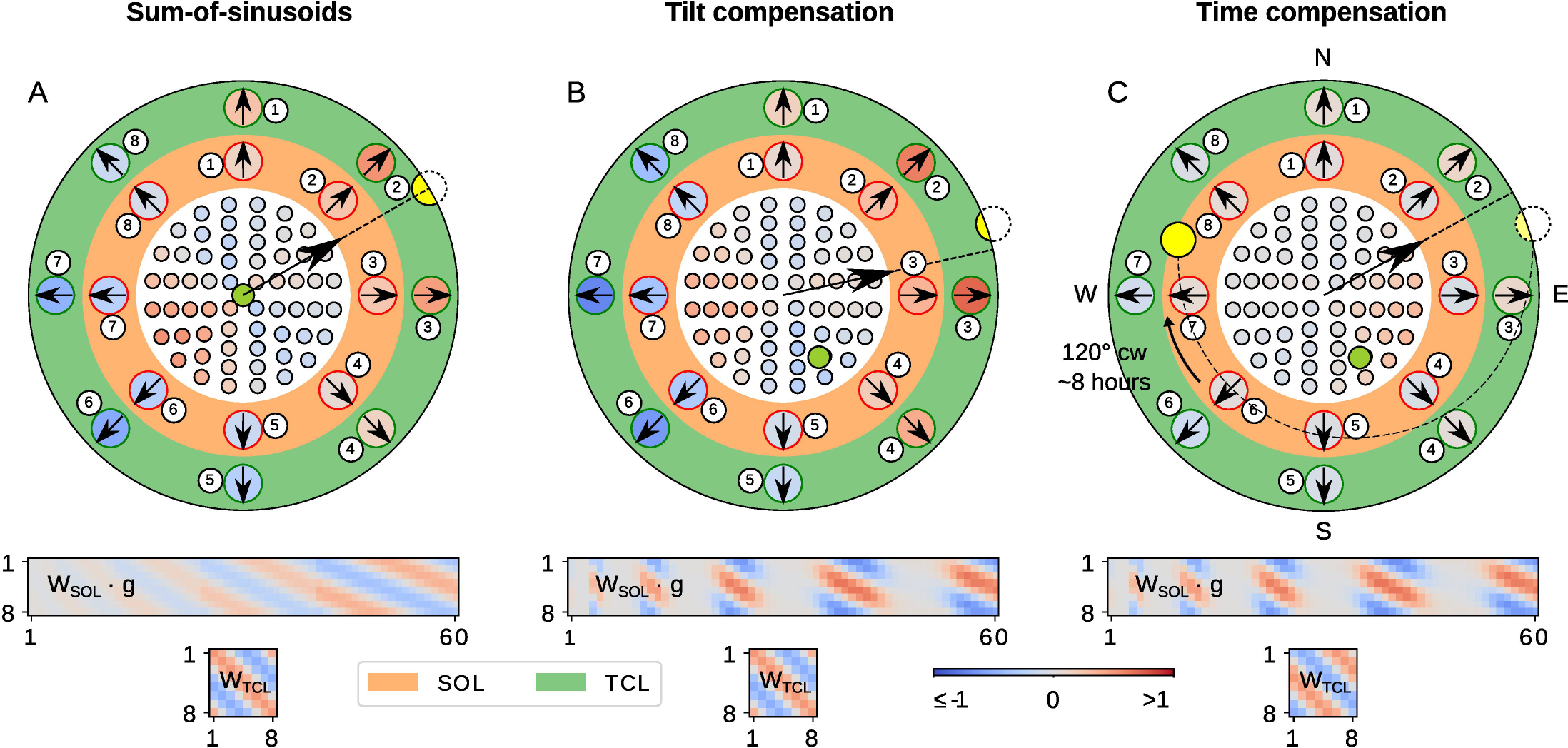
Step-by-step processing of the compass model. The white, orange and green areas show the response of the POL-, SOL- and TCL-neurons respectively; red denotes excitation and blue suppression (see colour-bar for values). The set of synaptic weights connecting each layer is shown under the disc (values based on the same colour-bar). Orange background means that the activity is affected by the gating function, and green that it is affected by the time compensation mechanism. In the white disc, different points are the relative positions of the POL units on the sensor; the numbering starts from the centre and unwraps clockwise towards the outline like a spiral (not shown); the round green mark is the point of the sensor that is aligned with the zenith of the sky, and the yellow circle is the sun position. The black arrow with the dashed line is the decoded prediction of the solar azimuth from the TCL-neurons; the numbering refers to the identities of the neurons in weight matrices below. The weight matrices show the synaptic weight between consecutive layers [defined by Eq. (6) and Eq (7) for the SOL and TCL respectively; values based on the same colour-bar]; the horizontal is the input and the vertical the output axis. (A) The sum-of-sinusoids mechanism detects the solar azimuth; the zenith (green) point is aligned with the sensor orientation; the solar azimuth is encoded in both the SOL- and TCL-layers and the activation code (i.e. phase) of the two layers look identical, as the time compensation mechanism has been deactivated. (B) The tilt compensation mechanism corrects the predicted solar azimuth using tilting information; the sensor has been tilted 30° NNW (so now the zenith point is 30° SSE); the gating function has changed the focus on the specific ommatidia, as shown in the **W**_SOL_⋅**g** matrix. (C) The time compensation mechanism corrects for the solar azimuth changes using the solar elevation; 8 hours have passed so the sun has moved 120° clockwise but the compass is still aligned to the same direction due to the updated **W**_TCL_ weights (see also the difference in SOL and TCL responses compare to previous steps).

## Evaluation

### Measuring compass accuracy

In order to evaluate the performance of the model, we introduce an objective function, *J*, which measures the average error across multiple predictions of the solar azimuth. Each prediction is made for a different sun position and tilting orientation. More specifically, we tested 17 tilting orientations, namely 8 homogeneously distributed on a 60° tilted ring, 8 on a 30° ring and one pointing towards the zenith. For each of those tilting orientations, we sample 500 homogeneously distributed sun positions on the sky dome, giving a total of 8500 predictions. The error is given by the following equation,

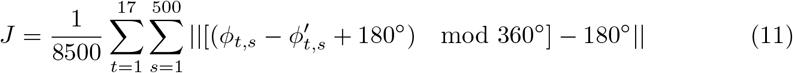

where *ϕ*_*t,s*_ and 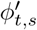 are the real and estimated solar azimuths.

We report the error as the mean absolute angular error plus/minus the standard error (MAE ± SE). A more detailed description of the objective function can be found in S3 Appendix. We also report the confidence of the predictions, *τ*_*s*_, as a value that shows how much we should rely on the respective predictions. Fig 8A gives a schematic representation of the objective function.

**Fig 8.**
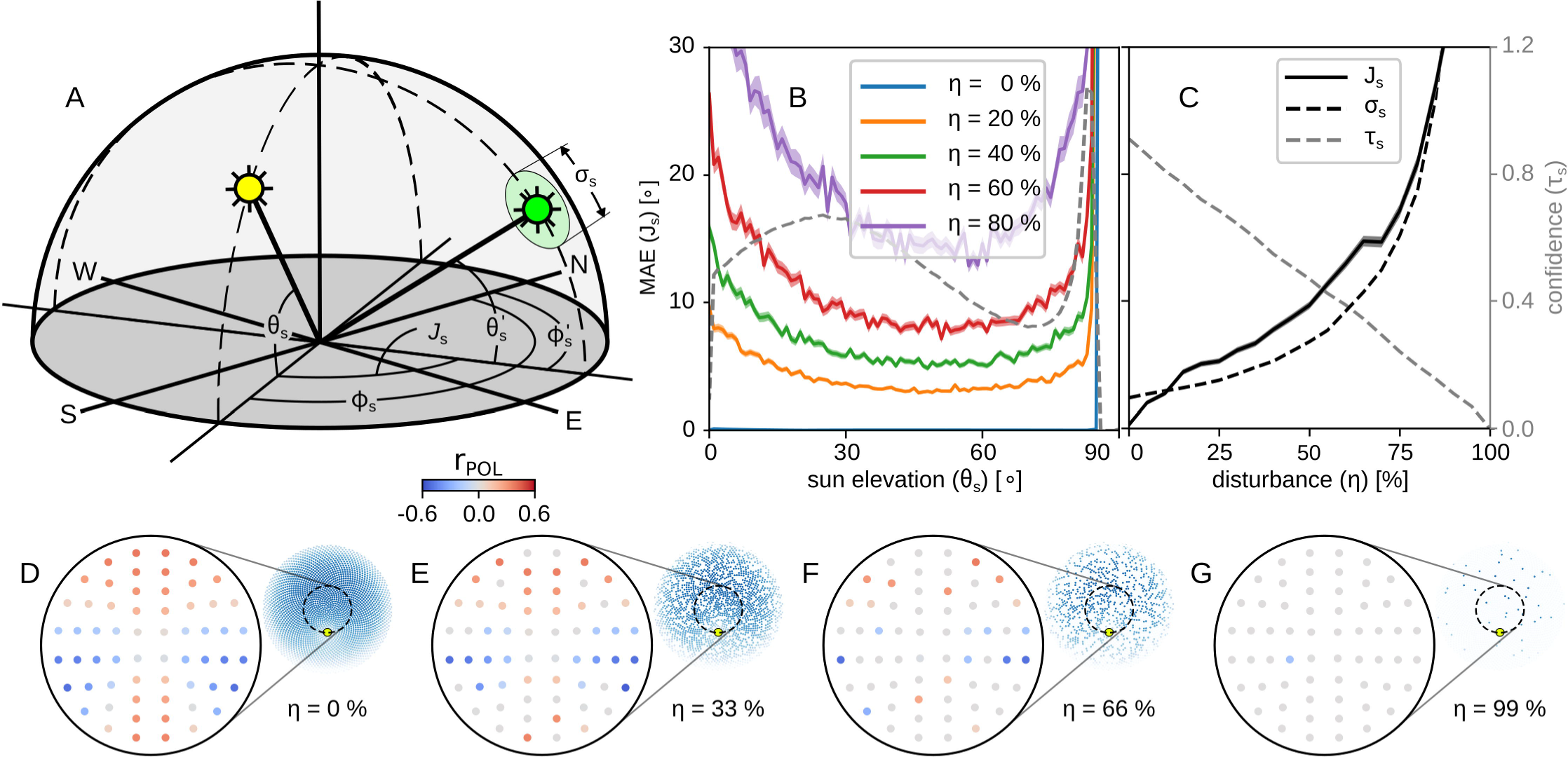
The objective function and the accuracy of the compass. (A) Schematic representation of the objective function; the yellow and green suns illustrate the real and estimated sun position; the disk around the green sun denotes the standard deviation of the estimation, 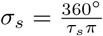. (B) Mean absolute angular error (coloured solid lines), MAE ±SE, and confidence (black dashed line), *τ*_*s*_, against the solar elevation for different disturbance levels, when the sensor points towards the zenith. (C) The mean absolute angular error (black solid line),standard deviation (*σ*_*s*_; black dashed line) and confidence (*τ*_*s*_; grey dashed line) against different disturbance levels. On the bottom there are some examples of the responses of the POL units in (D) a non-disturbed condition; (E) with *η* = 33 % disturbance; (F) with *η* = 66 % disturbance; and (G) with *η* = 99 % disturbance. The insets show a sample of the sky that caused the responses and the yellow mark on it shows the position of the sun.

The same objective function was used to explore the parameter space in both the design and the computational model of the sensor. The parameters for the layout of our sensor along with those of the computational model are inspired by biological features in insects, and mainly the *Cataglyphis* desert ants (see Table 1). Nevertheless, we are interested in exploring different set-ups that may perform better than the one that biology indicates.

### Sensory input disturbance

By adding perturbations of the polarisation signal in the simulation, we can evaluate the robustness of the sensor in noisy environments. In a natural environment, this could be caused by a sensor malfunction, clouds, vegetation or other objects that can block the light from the sky or destroy the polarisation pattern (note however that in the UV spectrum, clouds have a relatively small effect on light propagation). In practice, a disturbance level, 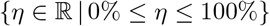, specifies the percentage of ommatidia that fail to contribute to the input. These are uniformly distributed across the surface of the eye. We tried different types of disturbance (including Gaussian noise) but found the type of disturbance shown here to be the most informative.

### Behavioural simulations

The aim of our model was to fill in the missing link between the compass information available from the sky and the path integration behaviour of insects. We therefore assess the performance using a simulated ant in a simulated world using the accurate sky model. The agent takes a predefined foraging path sampled from real ant data [54], which is translated into a sequence of directions and distances that the agent follows for its outward journey. It uses the insect eye and visual processing model described above as input to the CX path integration model [35], and the consequent CX output to control its inward journey. The ability of the agent to return home by the direct path is assessed using the same methods as in [35] allowing direct comparison between the use of simulated input and our network. We also test with conditions of additional sensor disturbance and head-tilt caused by uneven terrain.

### Simulating neurophysiological experiments

As outlined in the introduction, neurophysiological recordings from the protocerebral bridge of the desert locust [33] appears to show a compass output that spans only [0°, 180°]. To see if we can account for this result, which is discrepant with our model, we perform a simulated neurophysiology study in which we record the response of the TCL-neurons in our model using the same stimulus conditions as the animal experiments. Thus, we expose the artificial DRA to a uniform light source filtered by rotating linear polariser, and construct tuning preference curves for the TCL-neurons.

## Results

### Compass accuracy without tilt

We first evaluate our compass model in conditions where it always points towards the zenith. The average error in the absence of disturbance is *J* = 0.28° ± 0.1620° for *N* = 1000 sun positions homogeneously distributed on the sky-dome, and the average confidence was *τ*_*s*_ = 0.91, which is quite high (see Fig 8B and C to compare with regular confidence levels).

The solar elevation strongly affects the polarisation pattern in the sky, and as a result the accuracy of the compass predictions. Fig 8B reports the error measured for different solar elevations, and different levels of disturbance. Note we measure the elevation as the angular distance from the horizon (thus the zenith corresponds to an elevation of 90°). The blue line, which is hardly visible and lying on the bottom of the figure, is the error for samples without disturbance. The dashed line is the average confidence reported across all the reported disturbance levels. It is not hard to notice that, apart from the error, the confidence is also affected by the solar elevation.

In the model, we use this latter effect to our advantage, to estimate the elevation from the confidence. Fig 9A-C show how the different disturbance levels affect the estimation of the solar elevation,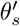 (black dots – real, red dots – predicted), using the confidence, *τ*_*s*_. Fig 9D shows the predicted against the real solar azimuth change rate.

**Fig 9.**
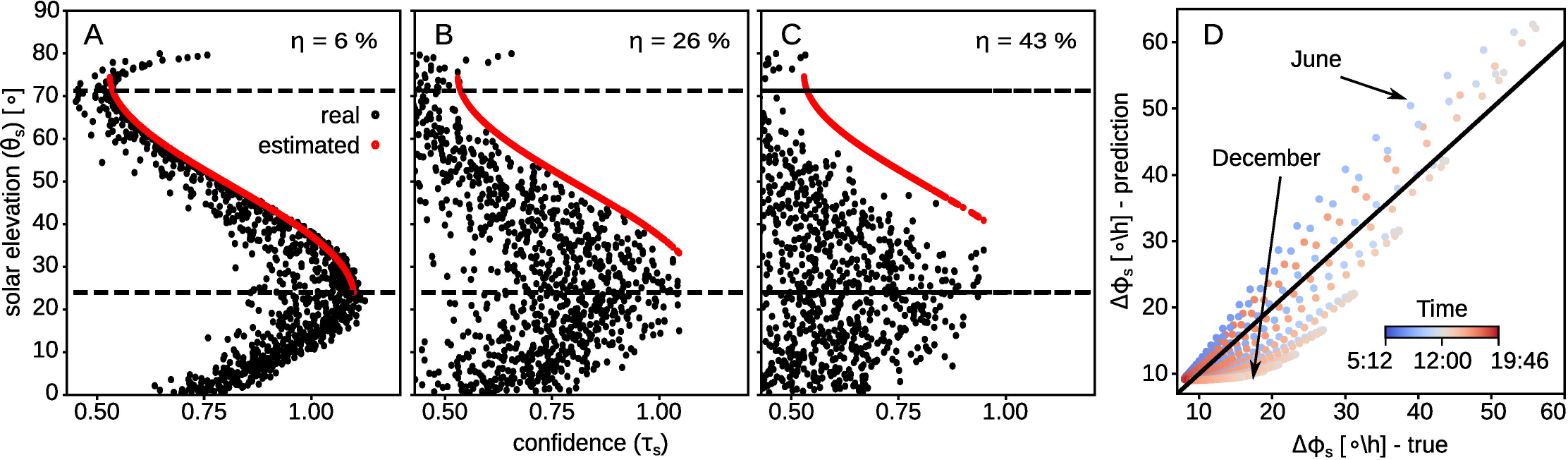
Dealing with time and light disturbance. Transformation of the compass response to solar elevation, and to the derivative of the solar azimuth function. (A - C) Function of the elevation with respect to the response with disturbance *η* = 6%, *η* = 26% and *η* = 43%; the red dots are the predicted solar elevation given the compass response, and the black dot denote the real values; the dashed lines give the range of the function. (D) Function of the real derivative of the solar azimuth against its prediction using Eq (9); black line shows perfect match of the two derivatives; colour denotes the time: blue is for morning and red is for evening.

Fig 8C gives a summary of the effect of the disturbance on our model’s predictions. We notice that as the disturbance grows so does the error of the predictions, but the confidence drops. This suggests we should not trust the predictions of our model when the disturbance of the sensory input is more than 85%, but for lower disturbance levels the compass still gives predictions with less than 30° error, which can be sufficient for the path integration task (see later results).

#### Effects of head tilt

Navigating ants are subject to large changes in their head pitch angle, particularly when carrying objects such as food or nest mates [52]. Here we assess how this might impact the accuracy of their celestial compass.

As we described in the methods, we filter the connections between the POL- and SOL-neurons using a gating function. With this function deactivated, and thus all ommatidia providing input to the solar layer, the performance of the model drops significantly. Fig 10A, B and C demonstrates the increased error of the predictions for different sun positions as the sensor is tilted for *δ* ≈ 0°, 30° and 60° respectively, and Table 2 summarises the respective average error along with the overall error.

**Table 2.** Mean absolute error before and after using the gating function.

|  | before gating |  |  | after gating |  |  | N |
| --- | --- | --- | --- | --- | --- | --- | --- |
| $J_{\delta=0^\circ}$ | $0.63^\circ$ | ± | $0.01^\circ$ | $0.47^\circ$ | ± | $0.01^\circ$ | 500 |
| $J_{\delta=30^\circ}$ | $35.68^\circ$ | ± | $0.01^\circ$ | $9.53^\circ$ | ± | $< 0.01^\circ$ | 4,000 |
| $J_{\delta=60^\circ}$ | $104.01^\circ$ | ± | $0.01^\circ$ | $13.16^\circ$ | ± | $< 0.01^\circ$ | 4,000 |
| $J$ | $65.78^\circ$ | ± | $0.62^\circ$ | $10.47^\circ$ | ± | $0.12^\circ$ | 8,500 |

N, number of samples used for this error; $J_{\delta=0^\circ}$ , MAE ± SE for $\delta = 0^\circ$ tilting; $J_{\delta=30^\circ}$ , MAE ± SE for $\delta = 30^\circ$ tilting; $J_{\delta=60^\circ}$ , MAE ± SE for $\delta = 60^\circ$ tilting; $J$ , overall MAE ± SE; MAE, mean average azimuthal error; SE, standard error.

**Fig 10.**
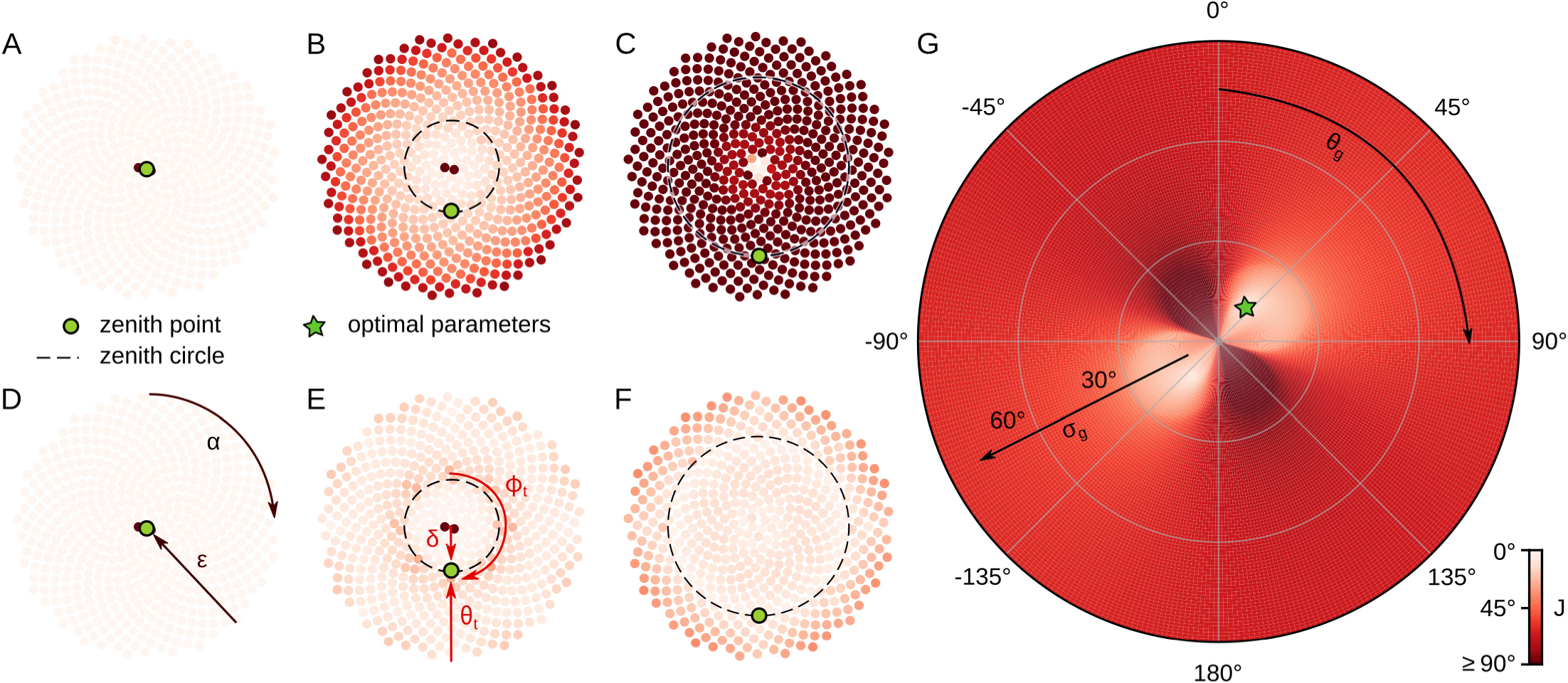
Dealing with tilt for a variety of gating parameters. Angular error of the expected direction of the sensor for different tilted angles and gating parameters. Black arrows show the axes; red shading shows the value of the objective function (*J*) - darker shading is for higher error values; red shaded points show the values of *J* for different sun positions; green discs show the zenith angle, *θ_t_*; black dashed lines visualise this angle for any tilt direction, *ϕ*_*t*_ (red arrows). (A-F) Visualisation of the azimuthal angular error with respect to the sun position, i.e. *ϵ* ∈ [0, 90] and *α* ∈ [0, 360), for three tilting angles of the sensor – (A, D) *δ* ≈ 0° (*θ*_*t*_ = 90°), (B, E) *δ* ≈ 30° (*θ*_*t*_ = 60°), (C,F) *δ* ≈ 60° (*θ*_*t*_ = 30°); without [top row (A-C)] and with gating [bottom row (D-F)]. (G) Average angular error for different gating parameters. The lowest cost (green star; *J* = 10.47° ± 0.12°, *N* = 8,500) is for ring radius *θ*_*g*_ = 40° and width (variance) *σ*_*g*_ = 13°.

Activating the gating function, the influence of each ommatidium to the responses of the solar layer becomes a function of the tilting parameters, producing much more robust results (Fig 10D-F). More specifically, the overall average error drops to *J* = 10.47° ± 0.12° (*N* = 8,500).

The default parameters for the gating function were selected using exhaustive global optimisation. More specifically, we fixed the design and network parameters and explore the different combinations of the parameters, *θ*_*g*_ and *σ*_*g*_, in the gating function. As the number of parameters in the function is very small and their range is also constrained, exhaustive global optimisation was not a very costly process. Fig 10G illustrates that the current combination of sensor layout and compass model perform best for a gating function with a ring shape of *θ*_*g*_ = 40° radius and *σ*_*g*_= 13° thickness.

The radius of the ring could be interpreted as the dominant angle of focus or the most informative direction. Moreover, our optimal main focus angle is 40° from the zenith point differs to the 25° angle, suggested in [25].

#### Exploration of the structural parameters

The sensor layout and the number of neurons in the computational model were based on biological data, but it is of interest to examine the effect of varying these parameters. The performance error of the compass (including tilted conditions) for a range of layout parameters is illustrated in Fig 11 A. The green line on this figure indicates the receptive field with the lowest error for the given number of units. The performance with respect to the receptive field seems to be independent from the number of units used. More specifically, the best performance on average was for *ω* = 55.99° ± 0.33°.

**Fig 11.**
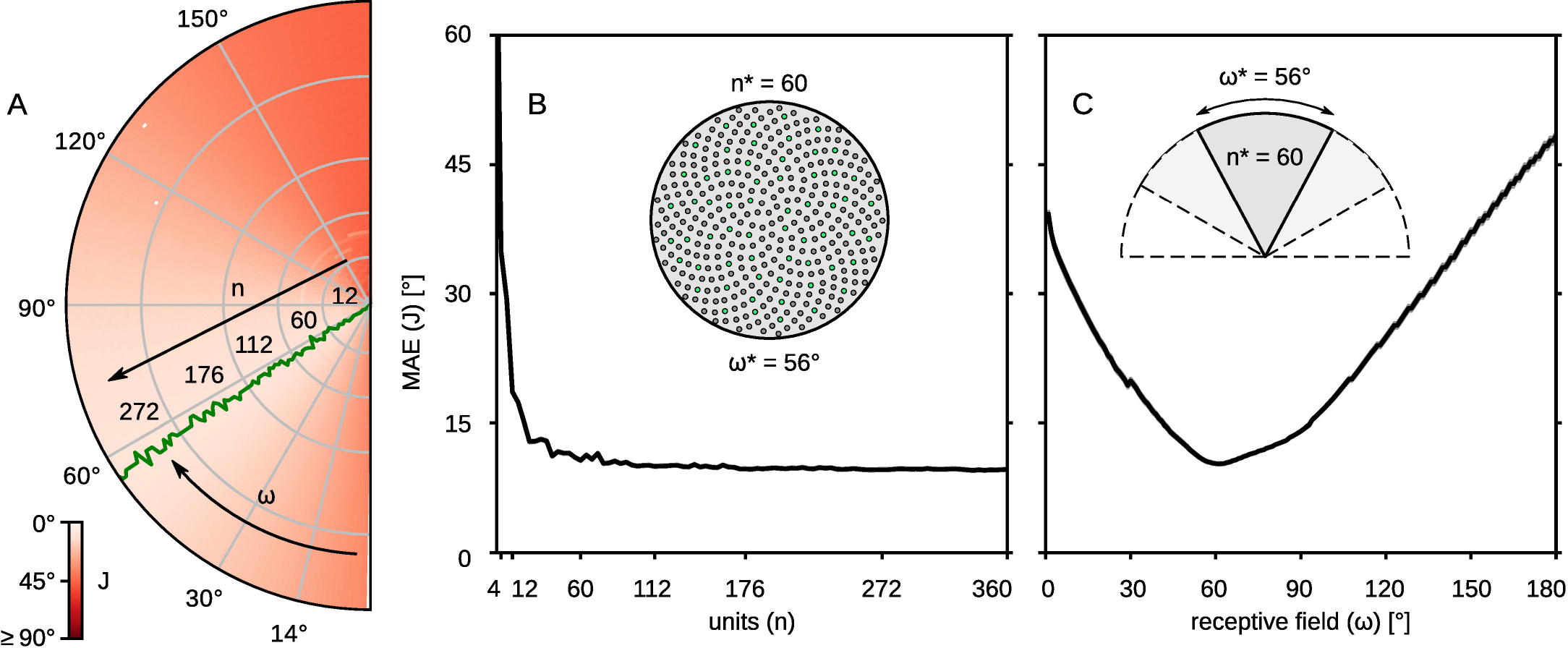
Optimal compass structural parameters. The performance of the compass for different topological parameters. (A) Values of the objective function on the *ω* × *n* plane; red shades illustrate the degree of error; black arrows show the axes; the green line shows the receptive field value associated with the minimum error for different number of units. (B) The error as a function of the number of units, *n*; the receptive field is fixed at *ω* = 56°; inset demonstrates the 56° wide sensor with *n* = 360 units; with green are marked the 60 units closest to the ones we chose. (C) The error as a function of the receptive field, *ω*; the resolution (ratio between *ω* and *n*) is fixed so that the number of units is *n* = 60 for *ω* = 56°; inset demonstrates the different visual fields including the optimal one with *n* = 60 units.

The error observed on the slice set by the green line in Fig 11A, where *ω* ≈ 56°, is illustrated in Fig 11B. This figure shows that there is a sharp drop of the error up to *n* = 60 units, after which there is not a significant improvement. A slice on the other axis, for *n* = 60, is illustrated in Fig 11C, which shows that the best design parameters for the sensor are *ω* ≈ 56° receptive field and *n* = 60 number of units. The average error reported for these parameters is *J* = 10.47° ± 0.12°. However, the lowest error reported was *J* = 9.55° ± 0.12° for *n* = 336 and *ω* = 56°.

The number of SOL- and TCL-neurons were selected based on the CX model described in [35]. However, we explored different populations of neurons and compare the performance to the one of the proposed model. Our results showed that less than 8 SOL inteneurons increase the error to *J* = 47.49° ± 0.32°, while more interneurons do not change the performance. Similarly, as long as we have at least 4 TCL-neurons, the performance of the sensor does not change for any number of SOL-neurons.

#### Path integration

To demonstrate the performance of the sensor in a more realistic scenario, we integrated the compass and path integration [35] models, testing how the compass accuracy affects the foraging and homing paths. We create an environment with a simulated sky and let an agent navigate in it. We guide our agent to food-source, using 133 different routes from Spanish desert ants *Cataglyphis velox* [54] and let the agent return to the nest using its path integrator. In addition, we test the performance of the agent under different sky conditions, by adding disturbance to the polarisation pattern.

Fig 12 summarises the results of the above experiment. The faded coloured lines in Fig 12A (even terrain) and B (uneven terrain, illustrated in Fig 12C) are the outward paths and the bold lines are the inward paths. The colour of the line identifies the different disturbance level. We use similar evaluation methods to [35] to allow direct comparison. Fig 12D and E show the overall performance of the agent in the path integration task with respect to the tortuosity of the inward route, 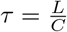, where *L* is the distance from the nest and *C* is the distance travelled.

**Fig 12.**
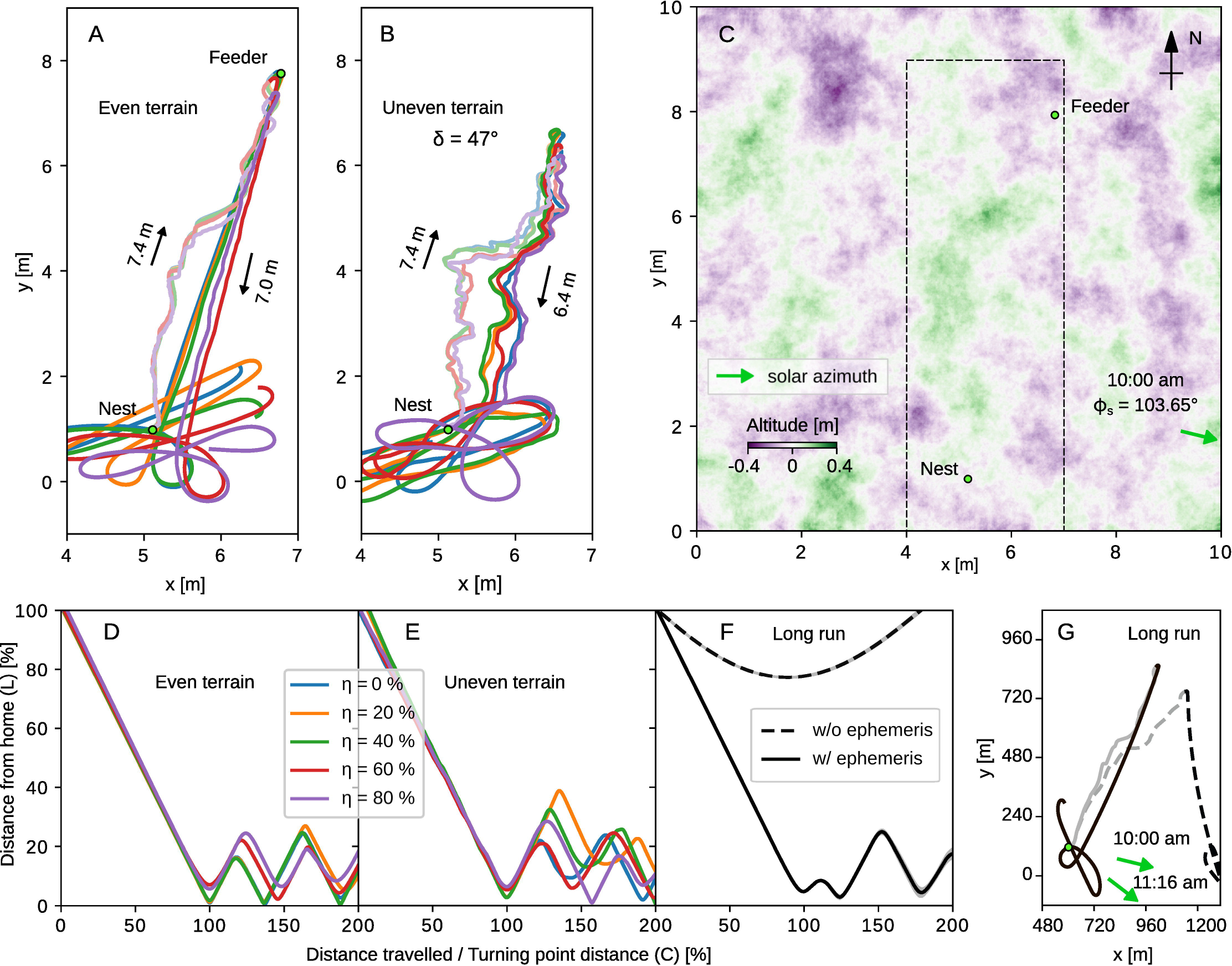
Behavioural simulation for the path integration task. Testing the celestial compass on path integration tasks. We set up the experiments to take place at 10am in Seville, Spain (37°23^′^33.03^′′^N, 5°53^′^01.95^′′^W). The altitude variance is 0.8 m and the maximum tilting angle noticed in all the experiments is *δ* = 47°. (A) Five representative routes of ants in different sky disturbance levels for an even [(B) uneven] terrain and their respective inward paths; different colours are for different disturbance levels (see legend); the faded lines are the outward paths and the bold ones are the inward. (C) The uneven terrain map; green colour denote hills and purple valleys; the marked region is the one cropped for the A and B plots. (D) Deviation from the best possible route during homing for different disturbance levels for even [(E) uneven] terrain. We scale up our experimental arena (by a factor of 120) to enable longer runs that demonstrate the performance of the time compensation mechanism. (F) Comparison of the path integration performance in terms of tortuosity with (solid black line) and without using the time compensation mechanism (dashed line). (G) The actual paths generated by the above experiment; green arrows show the direction of the sun at the beginning (10:00 am, 103.65° clockwise from north) and end of the route (11:16 am, 127.47° clockwise from north).

The results show that in most cases, the agent is moving in the correct direction until it reaches the nest and then does a systematic search for the nest. An exception is for *η* ≈ 97% (figure in S6 Fig), where the agent continues moving in the same direction not realising how far it has travelled. Overall, for disturbance *η* ≤ 90% (not shown in figure) the agent seems to navigate without noticeable problems. However, for higher disturbance levels we see a drop in the performance of the navigation task, walking at least twice the distance of the nest from the feeder for *η* ≈ 97%. The performance of the agent is affected very little by the uneven terrain, which shows that the tilting of the agent is not a problem anymore. The figure in S5 Fig summarises the results for different input disturbance levels and steepness of the terrain. Figure in S6 Fig illustrates the corresponding detailed paths of all the agents.

The terrain used here is drawn from a normal distribution, allowing the agent to tilt for a maximum of *δ* = 47°. The outward paths are consequently distorted by compass and distance errors while following the predefined sequence of directions and distances, but the homing paths still successfully guide the agent back to the nest, suggesting that any systematic bias in compass or distance information caused by uneven terrain is balanced out between the outward and inward routes. However, it is clear that the uneven terrain introduces an extra level of moment by moment disturbance in the heading direction.

In addition, we tested the performance of the sensor in longer runs, which take more time and hence will test the operation of the sensor’s time compensation mechanism (Fig 12F and G). We multiply the dimensions of the arena and the outward paths of the ants by a factor of 100, transforming the arena to 1 km × 1 km and the total run of the agent to 1 hour and 16 minutes. In this time (from 10: 00 am to 11: 16 am) the sun position changes by 23.82° clockwise. Fig 12F and G show that including the time compensation mechanism the agent successfully returns to the nest, while without it the path integration mechanism leads it away from the nest due to the change of the sun position. For detailed paths of multiple ant routes see figure in S7 Fig.

#### Experimental paradigm

We have noticed that the output of our compass model, the TCL-neurons, is not identical to the electrophysiological responses of the locusts’ TB1-neurons reported in [33]. However, this is not surprising, as the testing conditions of the two experiments were very different. More specifically, in the locust experiment, the animal was pinned in a vertical position and its DRA was exposed to uniform light passing through a rotating polariser. On the contrary, we expose our sensor to realistic sky-light facing upwards, assuming that the head of the hypothetical animal is aligned with the horizon. Therefore, we tried to simulate the former experimental environment and compare the responses of our TCL-neurons to the TB1-neurons recorded from the desert locust.

We found that the response of the simulated compass neurons closely resembles the double preference angles recorded in locust TB1-neurons [33] when stimulated by a rotating linear polariser under a uniform light source (Fig 13B and D). This contrasts dramatically with the response of the same simulated neurons when exposed to the natural skylight pattern (see Fig 13F). Moreover, calculating the preferred directions from the response of the simulated neurons under the linear polariser, for each column (Fig 13C), produces results rather comparable to the locust (Fig 13A). Note that for the linear polariser, the preferred direction has an inherent 180 degree ambiguity: for our simulation data (Fig 13C) we resolve this by taking the stronger of the two peaks (effectively a random choice as this difference results from noise); for the locust data (Fig 13A) we use the direction chosen in the original paper. These data have been interpreted in [33] as supporting a [0, ∼ 180°] ‘map’ of polarisation directions across the protocerebral bridge, increasing by ∼22.5° per column (see fitted line, Fig 13A). Our results suggest this effect may be a consequence of the experimental procedure rather than revealing the true directional preference - relative to the sky pattern - of these neurons, which may instead resemble Fig 13E.

**Fig 13.**
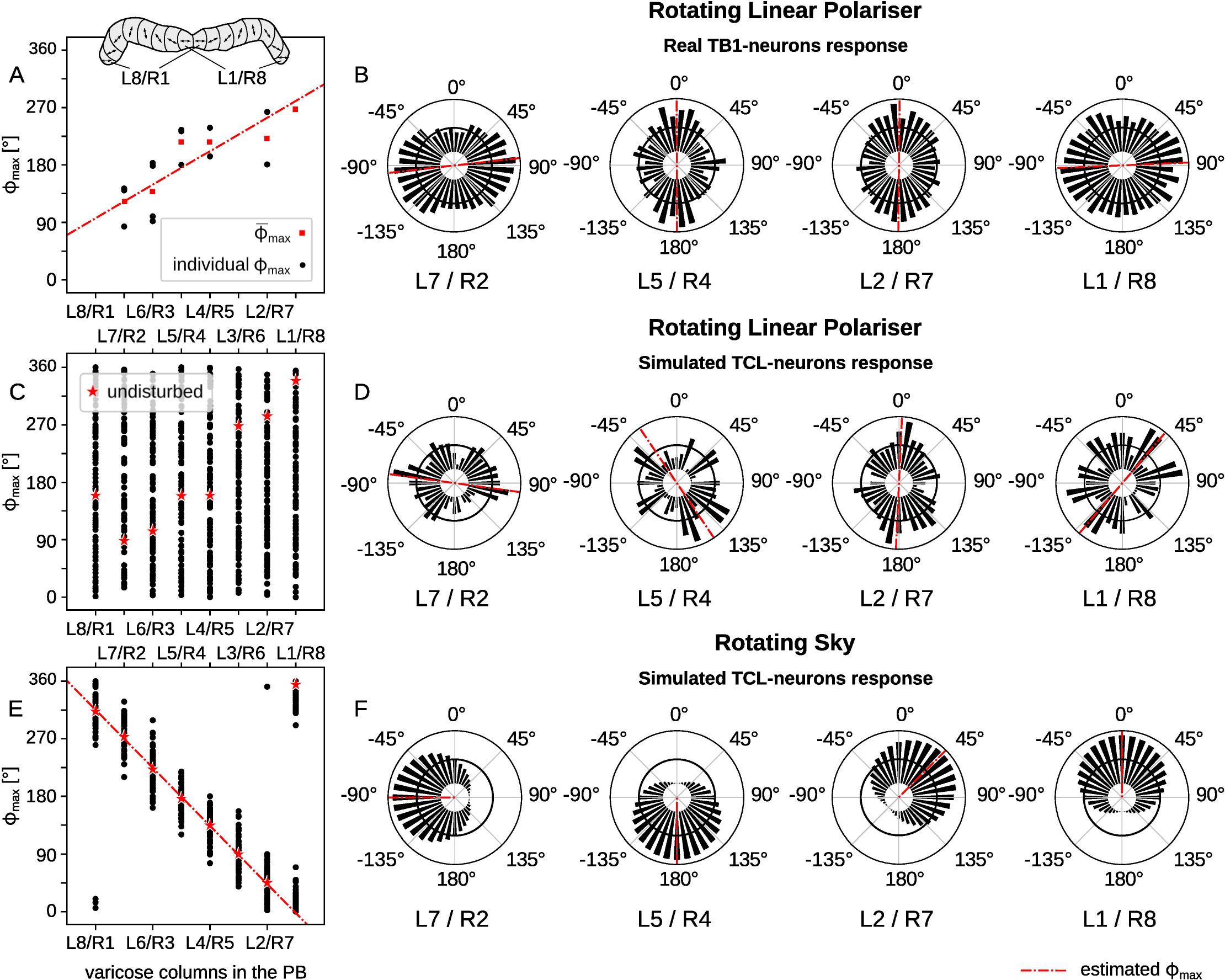
Real and simulated response of compass neurons for artificial and natural polarised light. E-vector orientation resulting in maximum excitation, *ϕ*_max_, for TB1-neurons in different columns of the CX (A,B) and TCL neurons in our simulation, determined by circular statistics (Rayleigh test [55]): (A) Real data of TB1-neurons (*N* = 15) in the PB of the locust brain, reconstructed from original data supplied by Stanley Heinze after [33] (inset shows the naming of the columns in the PB), (C) simulated TCL-neurons under a rotating linear polariser, and (E) under a rotating natural sky (*N* = 100, *η* = 50%, *θ*_*s*_ = 30°); black dots show individual samples, red squares show the mean *ϕ*_max_ and red stars show the *ϕ*_max_ in a condition without external disturbance (*η* = 0%); red lines are best linear fit to the data points. (B,D,F) show corresponding examples for specific neurons from four of the CX columns; bin width 10°; black solid lines indicate background activity; red lines indicate the *ϕ*_max_ direction.

The responses of Fig 13B and Fig 13D are similar but not identical, for example the TB1 recordings appear to show a 180° range of *ϕ*_max_ (Fig 13A) while our model gives a 270° range (Fig 13C). However, the number of recordings from the locust brain is limited and shows substantial variability (see the full set of simulation and locust responses in S8 Fig). As a consequence it does not seem productive to attempt to quantitatively replicate the details of this activity pattern with our model, but rather would be more interesting to test the response of these neurons to a more realistic sky pattern, which we predict should have a substantial qualitative effect on the observed activation patterns.

## Discussion

To perform path integration, insects need to transform polarised skylight to a global orientation. We have proposed a mechanism to explain how this might occur in the insect brain, given known anatomical constraints of the optic lobe and the protocerebral bridge. Our optic lobe model contains a large number (∼60) of polarisation-opponent neurons that respond to the degree and direction of polarisation received from each ommatidium. The protocerebral bridge should (to match previous work on path integration) contain exactly 8 compass neurons with sinusoidal tuning to 8 cardinal directions, ideally able to compensate for tilt and time. We show that this can be obtained by a weighting function that takes into account the fact that each ommatidium points at a specific region of the sky, and that the sky polarisation pattern has a specific relationship to the sun position. Most previous models have either assumed a simple linear polarisation as input [45], or, if physically implemented and tested under real skylight, have used a limited number of receptors (for a review see [56]). However, neurophysiological investigation of the receptive fields of central complex neurons suggests they could act as matched filters to the specific pattern of polarised light in natural skylight [29]. By combining a realistic sky model and receptor layout we show it is possible to obtain a good estimate of heading direction, sufficient for accurate path integration, robust to disturbances, able to compensate for head tilt that might occur with rough terrain, and to adapt to sun movement, providing a powerful celestial compass sensor based on the polarisation pattern.

### Obtaining solar azimuth from polarisation information

In our model, each neuron in the SOL layer integrates the signal from all sensory units. The relative azimuth of the sensor unit in the array to the preferred direction of the compass unit determines its weighting. As a result, the response across the compass units effectively represents the direction in the sky with the highest degree of polarisation, the cross-solar azimuth, from which the solar azimuth can be directly inferred. Our model thus counters the common assumption that a polarisation-based compass sensor must inherently have a 180° degree ambiguity and requires some additional signal to resolve the 360° directionality (see e.g. [49, 57]). Another consequence is that the best compass performance is not when the sun is on the horizon (thus producing the maximum degree of polarisation, largely in one direction, in the zenith) as has been sometimes assumed. In fact, for sun exactly on the horizon or exactly on the zenith, precise symmetry in the resulting polarisation pattern will result in ambiguity and low confidence in our model. The highest confidence occurs when the cross-solar azimuth falls within the receptive field of the sensor (modulated by the gating function), which corresponds to a solar elevation *θ*_*s*_ ≈ 28° (i.e., nearer to, but not on, the horizon).

Note however that for higher elevations, specifically, for 60° ≤ *θ_s_* ≤ 90° the sun itself would fall in the receptive field of the sensor. This suggests a parallel processing system based on sky luminance could form a complementary mechanism for determining the solar azimuth that would be accurate for solar elevations where the polarisation one is not. A speculative pathway for this could originate in the two out of the eight photo-receptors in the ommatidia of the desert ants that are sensitive in a wider range of the spectrum [58] (see Fig 3C), which could detect a sufficient light intensity gradient, and (in a similar way to our POL to SOL processing) form a novel skylight intensity compass. Alternatively or in parallel, the position of the sun is likely to also be detected by the non-DRA ommatidia of the compound eye, which are much better equipped for this task due to their smaller acceptance angle [38]. The two pathways could then be combined in the CX to form a complete insect celestial compass, consistent with the observation [58, 59] that the insect’s compass appears to integrate multiple modalities of light [60, 61]. Specifically, recent neurophysiological results show that all polarisation sensitive neuron types in the CX also show azimuth-dependent responses to an unpolarised UV or green light spot [57, 62].

### Neurobiological plausibility

Our model represents a proposed mapping from POL to TB1 neurons in a computational form, i.e., using a weight matrix derived from theoretical considerations rather than following details of the neural connectivity in the insect brain. Here we consider whether there is a plausible neural substrate for this computation.

Tangential neurons (TL) of the lower central body (CBL) represent the actual input of the polarisation pathway to the CX, with at least three TL types which are all polarisation sensitive. As their name suggests, these neurons provide input tangentially across all columns in the CBL. As such, they could form the basis for the ‘fully connected’ mapping in our model between POL- and SOL-neurons, as CX columnar neurons innervating the same CBL region could potentially sample from all POL inputs. A plausible candidate would be the CL1 neurons: their receptive filed is about 60° wide and their signal-to-noise ratio is relatively low [11]. There is evidence that CL1-neurons may be homologous to the E-PG neurons of the ellipsoid body (EB) in flies [34, 63], which represent landmark orientation and are used for visual navigation. This suggests the same neurons may get information from the visual field, such as the horizon line, which could provide the pitch and roll information that our model assumes will be integrated at this stage of processing to correct for tilting. However there is no direct evidence as yet to support the existence of our proposed gating function. A fascinating possibility is that this putative TL-CL1 mapping, which would need a rather well-tuned set of weights to extract the compass heading, could be a self-organising network, or at least could be calibrated by the experience of the animal, e.g., if it makes coordinated rotations under the sky, as many insects have been observed to do [15, 64].

Columnar (CL1) neurons innervate the protocerebral bridge (PB) and are presumed to connect to TB1 neurons, hence this could form the physiological basis of our model’s connections from SOL-to TCL-neurons. In other insects, e.g. fruit flies, the equivalent neurons to CL1 have been shown to form part of a ring-attractor network to encode heading relative to a visual target [34], which can also hold and update this information in the dark (based on self movement). The PB has recurrent connections to the lower central body (ellipsoid body in flies) which could provide the feedback hypothesised in our model to compensate for time [65, 66]. Alternatively, it has been noted [67] that there is a potential neural pathway from circadian pacemaker circuitry in the accessory medulla to the PB, which could provide an alternative way to adjust the compass with the time of day. As shown in our results (see also [29]) it may be difficult to interpret the real encoding principles of these neurons using isolated cues if they have evolved to be tuned to the combined input pattern of the real sky, and are potentially modulated by time and the tilting orientation of the animal. Specifically, we see that the robust [0, ∼ 360°] representation of direction in our simulated TCL neurons appears to be a noisy [0, ∼ 180°] representation when using a rotating polariser as the stimulus, resembling the data from [33].

### The sensor array

We found that the resolution and receptive field of the sensor are optimal for *n* = 60 units resolution and *ω* = 56° receptive field. However, the DRA of *Cataglyphis* has more units (ommatidia) and broader receptive field in the frontocaudal axis (*ω_a_* ≈ 120°; see Fig 3A). This asymmetric design of the DRA might be explained if we assume the ant’s head is tilted more often around the mediolateral axis; as they do not appear to significantly stabilise their head orientation while running up and down hills [68] or while carrying a load [52]; whereas there is some evidence that they do stabilise when their body is tilted around the frontocaudal axis [68, 69]. Therefore more samples on the frontocaudal axis would increase the confidence of their compass when running. Other insect species have distinctive differences in the layout of their dorsal rim: in the precise number of ommatidia, their alignment pattern, their acceptance angle and their spectral sensitives (see Table 3; for a review see [56, 70]). These might suggest similar adaptations to the specific requirements of habitat, foraging time, task and motor control.

**Table 3.** Properties of dorsal rim ommatidia in different species.

|  | compass | ant [13, 36] | cricket [58, 71, 72] | bee [9] |
| --- | --- | --- | --- | --- |
| $n$ | 60 | $\sim 112$ | 530 – 680 | $\sim 120$ |
| $\rho_{\max}$ | $5.4^\circ$ | $5.4^\circ$ | $35^\circ$ | $14^\circ$ |
| $\lambda_{\max}$ | 350 nm (UV) | 350 nm (UV) | 440 nm (blue) | 350 nm (UV) |

$n$ , number of facets; $\rho_{\max}$ , acceptance angle; $\lambda_{\max}$ , spectral sensitivity.

### POL-compass design for robotics

In this study we used a somewhat generalised DRA, as it was specifically our intent to consider how we might construct an equivalent sensor for a robot, preferably at a low cost. The insect celestial compass has already inspired the design of a number of polarisation compass sensors, particularly as a robust alternative to magnetic compass sensing for robot applications [56]. The approach applied in the Sahabot [47, 49, 50] introduced a number of biomimetic aspects, including POL-OP units, but used only 3, each one with a relatively small acceptance angle, all pointing towards the zenith, and oriented in angles with 60° difference. The dominant polarisation direction was recovered either by rotational scanning or by using a look up table, with additional light sensing used to decide between the solar and cross-solar directions. The output angular error reported for flat terrain experiments is 1.5°. Chu et al. [73] improved this design, using blue filters on photo-receptors with wider acceptance angles, *ρ* ≈ 53°, to obtain a minimum of 0.2° angular error. Ma et al. [74] and Xian el al. [75] followed the same line, optimising the DOP and AOP extraction using the least-squares method. More recently Dupeyroux et al. [76, 77] used UV sensors with orthogonally aligned HNP’B polarisers that were rotated 360° every 42 seconds using a stepper motor. Calculating a compass direction from this method produced from 0.3° to 1.9° peak errors in clear and cloudy skies respectively. Similar robot implementations include those by [78, 79]. Alternative approaches use a camera [80–84] or multi-camera system [79, 80, 85], or specialised image sensors with different polarisation sensitivity for each pixel [56, 86]. Good results are obtained by Stürzl et al. [84], who built a single camera sensor with 3 lenses and 3 polarisers oriented in different angles. The sensor estimates the angle and degree of polarisation of the skylight, and fits a sky-model on the input stream to estimate the parameters, which are the solar azimuth and elevation. In addition, they also estimate a covariance matrix that shows the confidence of their prediction. Their approach also works in tilted environments by integrating inertial measurement unit (IMU) data. Finally, Yang et al. [87] follow up Sahabot’s work, use 2 POL-OP units oriented in different angles from the zenith, placed on a plane and use the scanning technique in order to get values from different angles. They show that their sensor can estimate the solar azimuth and elevation in clear sky conditions with MAE 0.2° and 0.4° respectively.

Our model suggests an alternative sensor design, in which a larger number of POL-OP sensors are used to sample specific areas of the sky, but these do not form a complete image as in the camera-based systems described above. Such a sensor could be built from off-the-shelf components, e.g., using pairs of UV photodiodes and linear polarisation filters to imitate dorsal rim ommatidia photo-receptor neurons. The components could be mounted on a dome, creating a similar DRA to ant eyes. As we have shown, the subsequently computation to recover heading direction is relatively low-cost and could easily be carried out on a robot-compatible microprocessor, which could also run our path integration model. We hope to build this sensor and test it on a robot in future work.

## Conclusion

As well as building a physical implementation, there are several other ways in which this model could be developed in the future. One consideration already discussed above would be to integrate parallel processing of luminance and spectral cues, which can provide complementary information to polarisation, and thus enhance the reliability with which the solar azimuth can be determined over a wider range of conditions. A second would be to examine whether a more direct mapping can be made between the computational processing we have proposed and the detailed neuroanatomy of the layers intervening between the POL neurons in the medulla and the compass neurons in the CX. Finally, we believe it is key that such a model remains grounded in understanding of the real task constraints the circuit needs to support, i.e., the natural environment conditions under which insect path integration evolved and operates.

## Acknowledgments

This work was supported by EPSRC grant EP/M008479/1 “Invisible cues”. We thank Stanley Heinze and Uwe Homberg for providing the biological data in Fig 13 and S8 Fig and discussion of these results. We also thank Timothy Warren and the other anonymous reviewers for their helpful feedback.

## Supporting information

**S1 Fig. Detailed view of the sensor layout.** Top and side view of the sensor layout, including all the design parameters: *n* = 60 units; *ω* = 56° receptive field; *ρ* = 5.4° acceptance angle for each ommatidium; ∆*ρ* = 5.6° interommatidia angle. The design was built using the Adobe Inventor tool and modified using the Inkscape.

**S2 Table. Spherical coordinates of the exact positions of the POL units on the dome.** ID (*i*), the identity of the POL unit referring to the figures of the main text; *θ*^*i*^, the zenith distance of the *i*^th^ unit; *ϕ^i^*, the azimuth of the *i*^th^ unit. The orientation of each unit is a function of its relative azimuth to the heading direction of the dome: *α*^*i*^ = *ϕ*_*i*_ − 90°.

**S3 Appendix. Details of the objective function.**

**S4 Appendix. Integrating the model for infinite SOL-neurons.**

**S5 Fig. Behavioural simulation for the path integration task - summarised results.** Outward (away from the nest – bold lines) and inward paths (towards the nest – faded lines) of artificial ants that use the proposed compass to orient themselves in different disturbance and inclination levels. Each panel of the top row shows the route of the ant in different maximum surface steepness (*δ* ∈ {15°, 28°, 38°, 47°, 52°}; different coloured lines in the same panel) which is associated with the respective altitude variance in the terrain shown in Fig 12C (*α* ∈ {0.2 m, 0.4 m, 0.6 m, 0.8 m, 1.0 m}; Fig 12 shows results for *α* = 0.8 m) for a specific sky-disturbance level (*η*). The panels of the bottom row show the routes of the ant in different sky-disturbance levels (*η* ∈ {0.0, 0.2, 0.4, 0.6, 0.8}; different coloured lines in the same panel) for a specific maximum inclination. The bottom further left panel shows the performance is case of no inclination at all (even terrain, *δ* = 0°). The outward paths were adapted from [54], and the inward paths are the outcome of the path integration network described in [35] using our approach to extract the TB1-neurons’ responses.

**S6 Fig. Behavioural simulation for the path integration task - detailed results.** Grid of panels showing the performance of our model when used for path integration for different sky-disturbance levels (different columns for different sky-disturbance levels) and maximum steepness of the terrain (different rows for different inclinations). Lines in each panel show the outward (away from the nest – red solid lines) and inward paths (towards the nest – black dashed lines) of 133 ants using the proposed compass to orient themselves. The figure shows that our model’s performance is not affected much by external disturbances (sky of steepness), as it is always able to return to the nest – with an exception for sky-disturbance of *η* = 0.97. Although the paths by themselves vary a lot due to the different inclinations, the ants are capable of successfully return to their starting points. The outward paths were adapted from [54], and the inward paths are the outcome of the path integration network described in [35] using our approach to extract the TB1-neurons’ responses.

**S7 Fig. Behavioural simulation for the path integration task - compensating for time.** Outward (away from the nest – red solid lines) and inward paths (towards the nest – black dashed lines) of artificial ants that use the proposed compass to orient themselves during their visit to a distant food sources of around 700 m away resulting in long runs. These runs take on average 1 hour and 16 minutes to complete and the solar azimuth changes for 23.82°, showing the effect of our time compensation mechanism. The outward paths were adapted from [54], and the inward paths are the outcome of the path integration network described in [35] using our approach to extract the TB1-neurons’ responses.

**S8 Fig. Real and simulated response of compass neurons - detailed results.** E-vector orientation resulting in maximum excitation, *ϕ*_max_, of the real (desert locust) and simulated (our model) neurons in the CX. Each panel shows the response (black bars) and *ϕ*_max_ (red line) of a specific TB1-(TCL-) neuron in the real (artificial) CX. The different rows of the panel-grid show the location of the neurons in the PB. The first three columns show the response of different TB1-neurons in the desert locust brain supplied by Stanley Heinze [33] and organised in rows with respect to their location; row 3 and 7 are empty due to a lack of data; different panels in the same row show different neurons in the same region. The fourth column shows the response of the artificial TCL-neurons in a similar stimuli to the one used in the first three columns. The last column shows the response of the same TCL-neurons when the artificial DRA is exposed to natural sky-light extracted by a simulated rotating sky.

