## Supplementary figures and images for "From skylight input to behavioural output: a computational model of the insect polarised light compass"

### S1 Fig

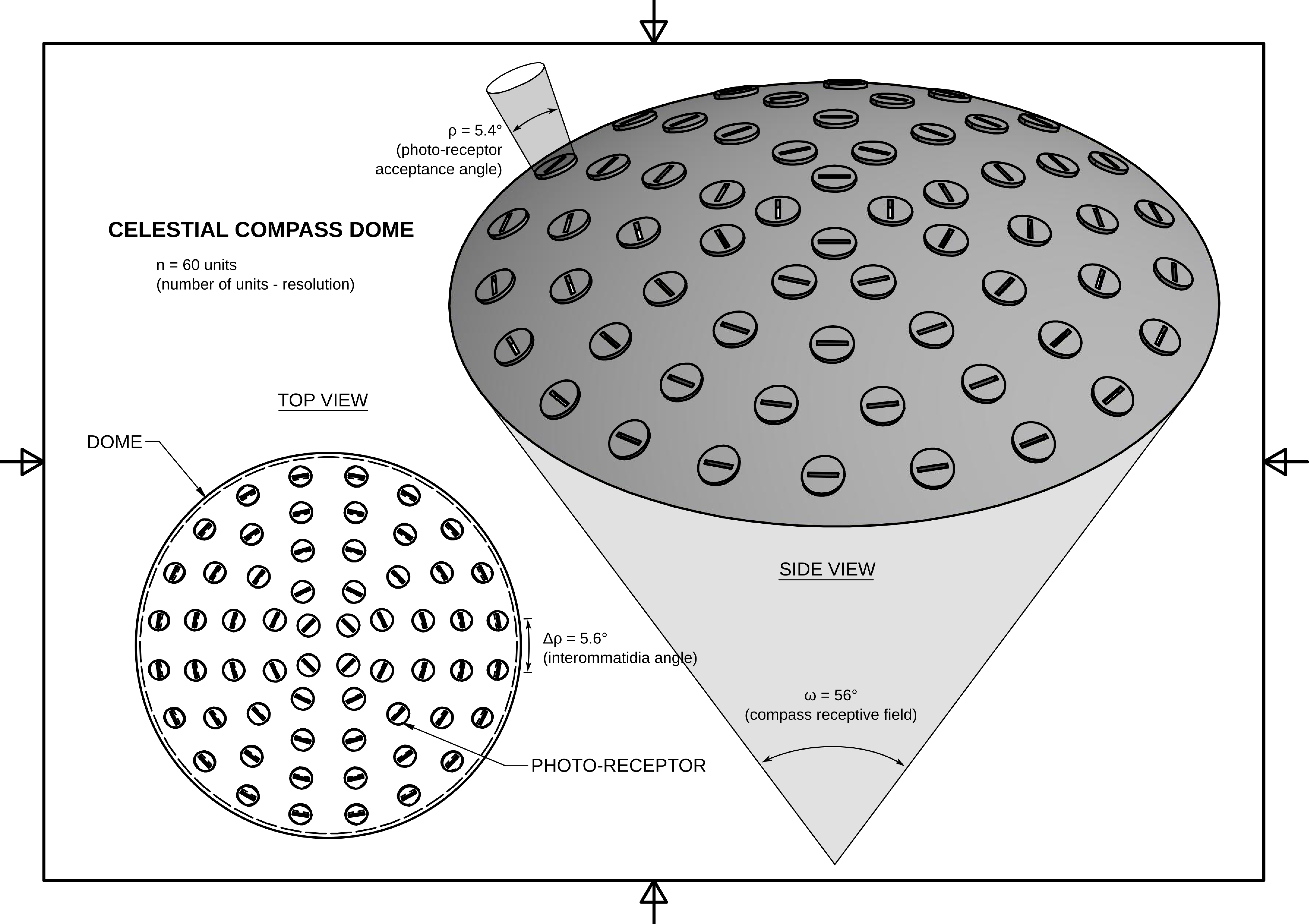

### S5 Fig

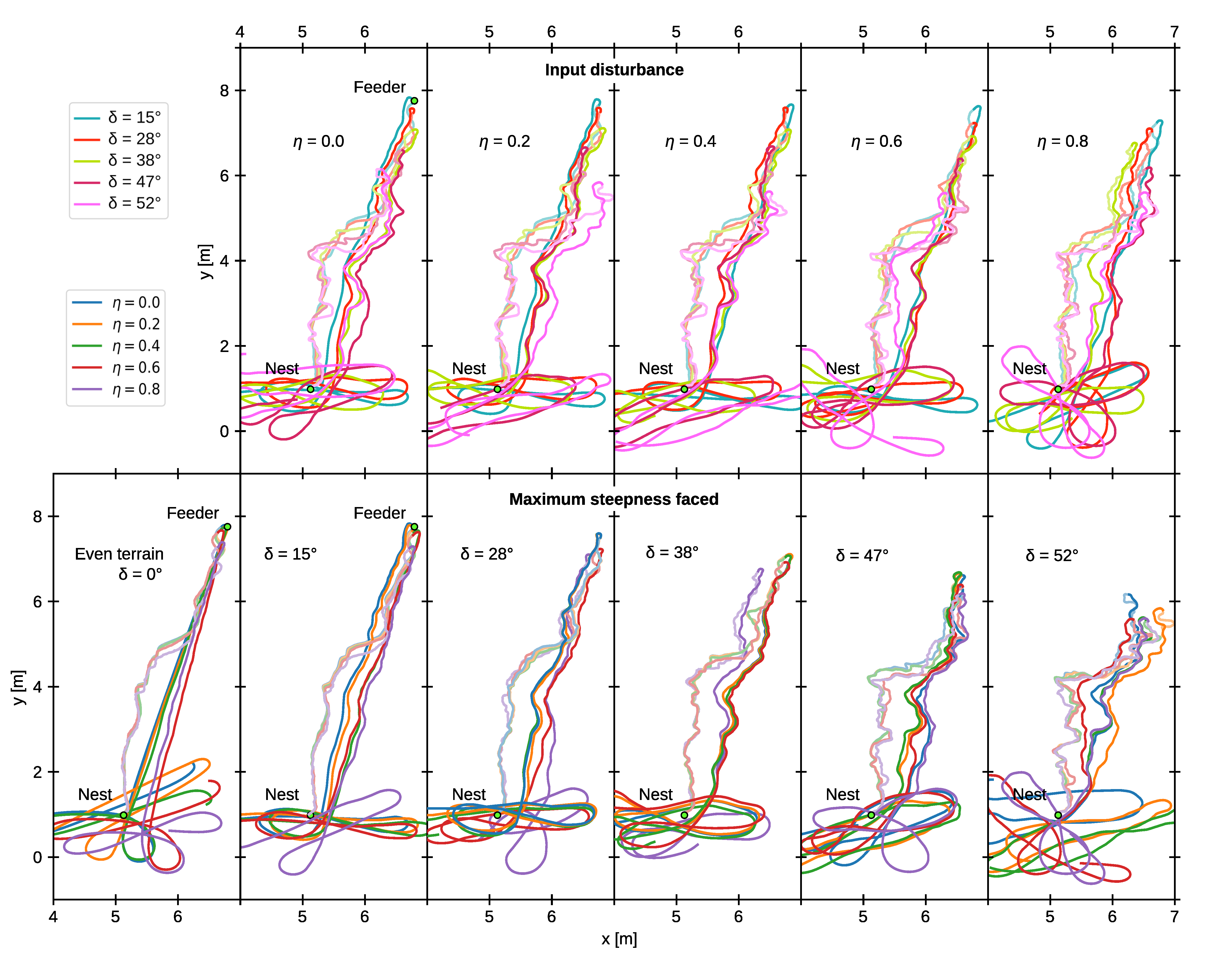

### S6 Fig

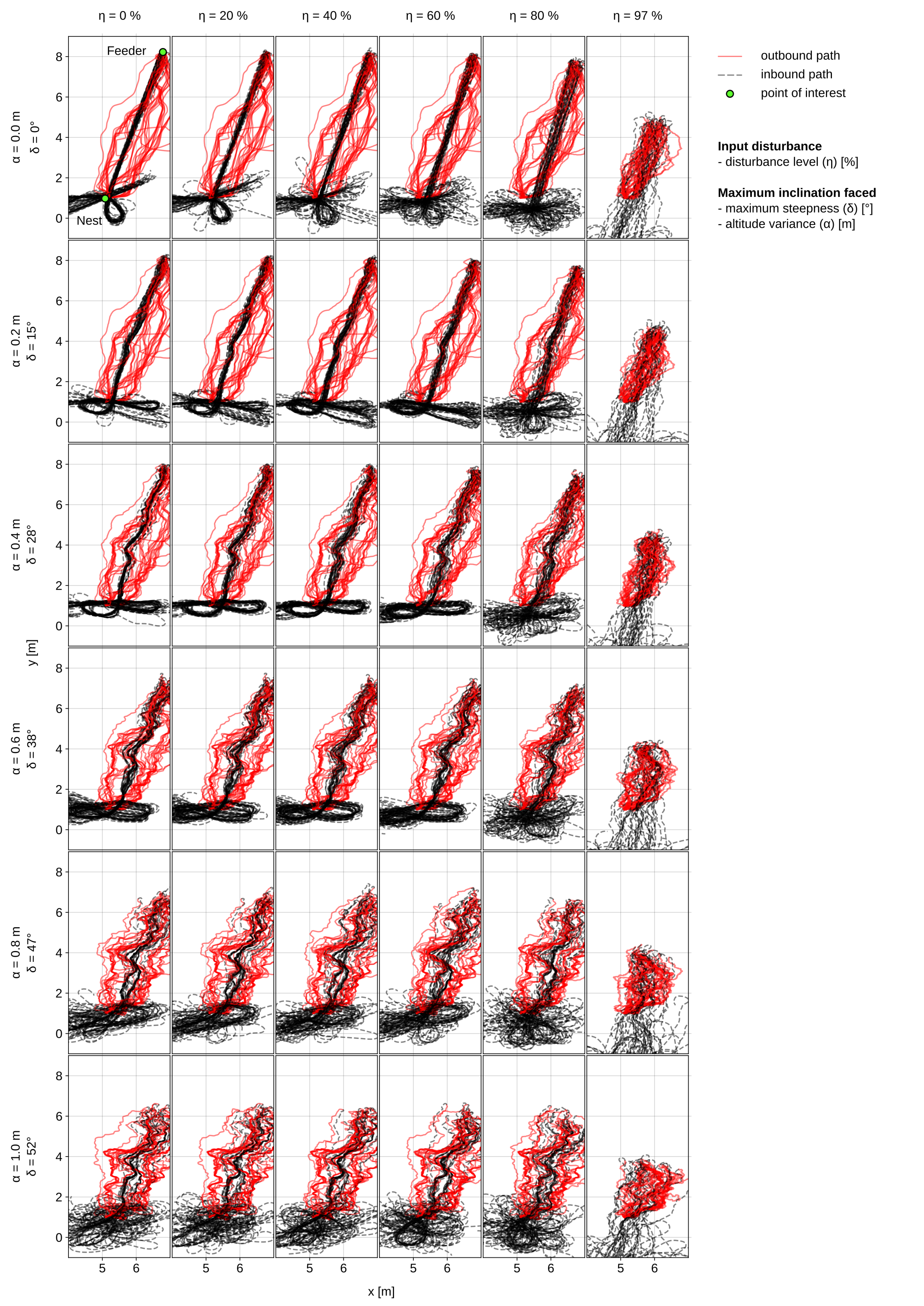

### S7 Fig

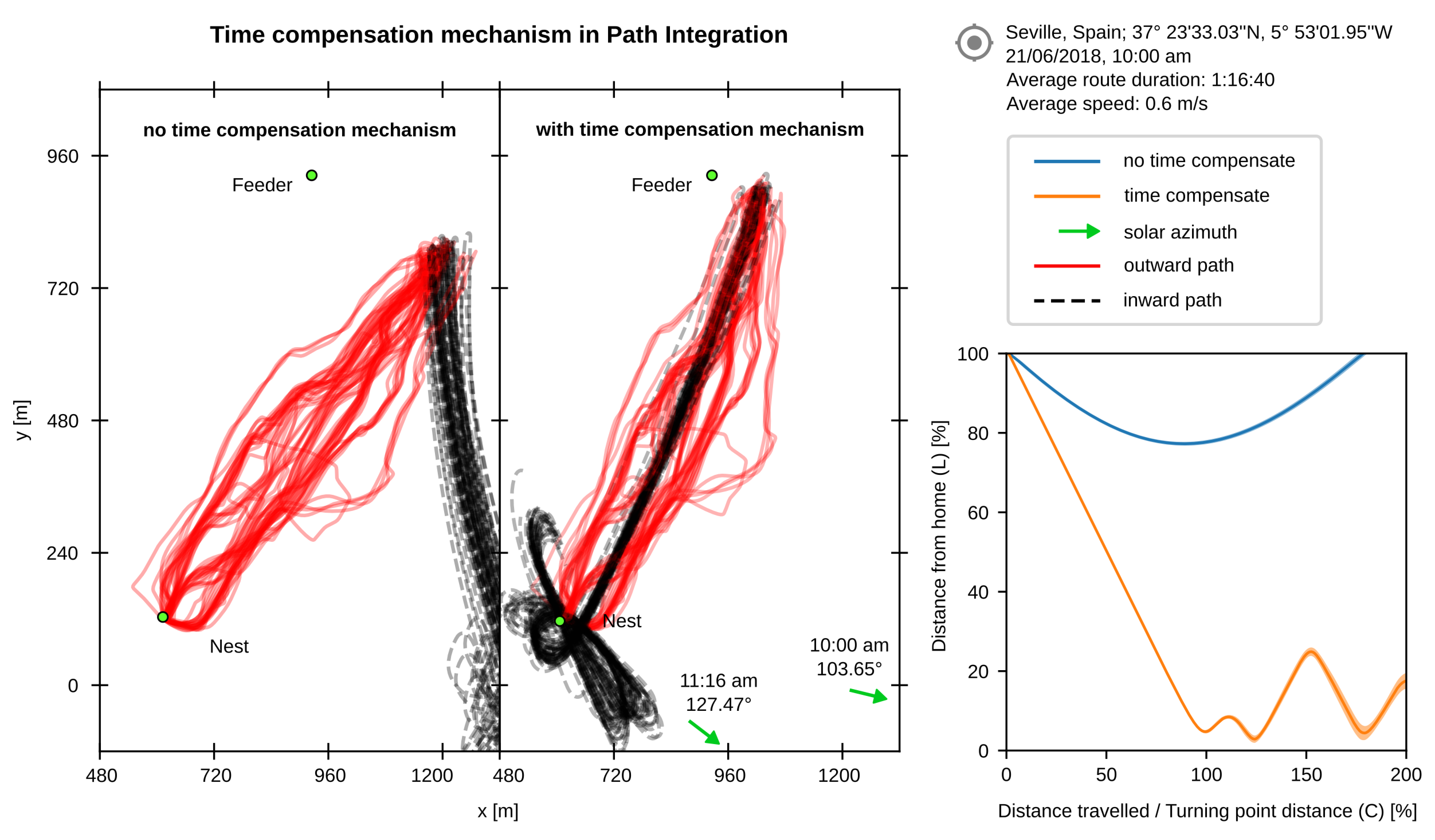

### S8 Fig

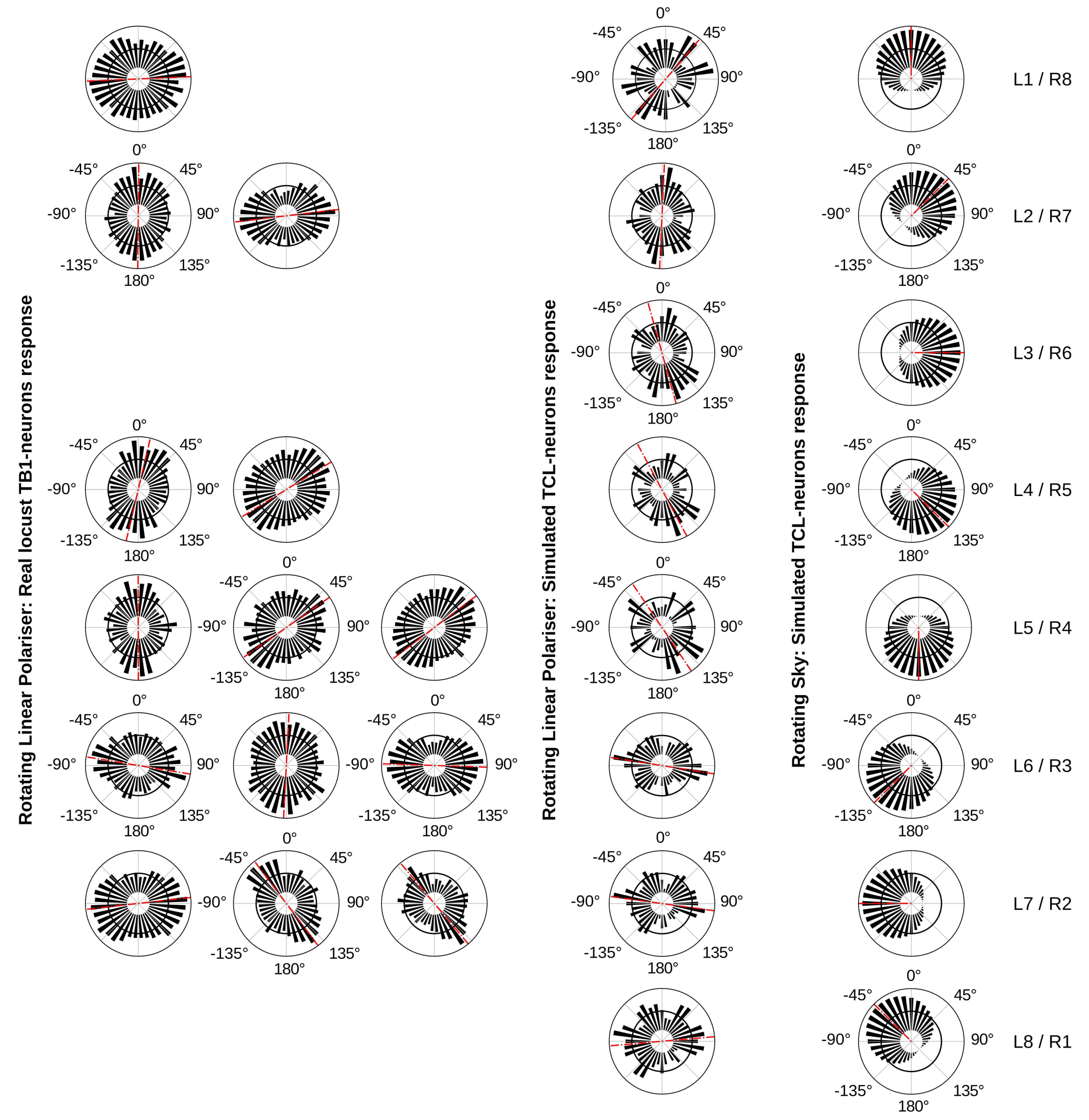
