## Supplementary material for "From skylight input to behavioural output: a computational model of the insect polarised light compass": S2 Table

**Table. Spherical coordinates of the exact positions of the POL units on the dome.**

| ID ($i$) | $\theta^i$ | $\phi^i$ | ID ($i$) | $\theta^i$ | $\phi^i$ | ID ($i$) | $\theta^i$ | $\phi^i$ |
|---|---|---|---|---|---|---|---|---|
| 1 | 5.13 | 45.00 | 21 | 15.39 | −105.00 | 41 | 25.65 | 9.00 |
| 2 | 5.13 | 135.00 | 22 | 15.39 | −75.00 | 42 | 25.65 | 27.00 |
| 3 | 5.13 | −135.00 | 23 | 15.39 | −45.00 | 43 | 25.65 | 45.00 |
| 4 | 5.13 | −45.00 | 24 | 15.39 | −15.00 | 44 | 25.65 | 63.00 |
| 5 | 10.26 | 22.50 | 25 | 20.52 | 11.25 | 45 | 25.65 | 81.00 |
| 6 | 10.26 | 67.50 | 26 | 20.52 | 33.75 | 46 | 25.65 | 99.00 |
| 7 | 10.26 | 112.50 | 27 | 20.52 | 56.25 | 47 | 25.65 | 117.00 |
| 8 | 10.26 | 157.50 | 28 | 20.52 | 78.75 | 48 | 25.65 | 135.00 |
| 9 | 10.26 | −157.50 | 29 | 20.52 | 101.25 | 49 | 25.65 | 153.00 |
| 10 | 10.26 | −112.50 | 30 | 20.52 | 123.75 | 50 | 25.65 | 171.00 |
| 11 | 10.26 | −67.50 | 31 | 20.52 | 146.25 | 51 | 25.65 | −171.00 |
| 12 | 10.26 | −22.50 | 32 | 20.52 | 168.75 | 52 | 25.65 | −153.00 |
| 13 | 15.39 | 15.00 | 33 | 20.52 | −168.75 | 53 | 25.65 | −135.00 |
| 14 | 15.39 | 45.00 | 34 | 20.52 | −146.25 | 54 | 25.65 | −117.00 |
| 15 | 15.39 | 75.00 | 35 | 20.52 | −123.75 | 55 | 25.65 | −99.00 |
| 16 | 15.39 | 105.00 | 36 | 20.52 | −101.25 | 56 | 25.65 | −81.00 |
| 17 | 15.39 | 135.00 | 37 | 20.52 | −78.75 | 57 | 25.65 | −63.00 |
| 18 | 15.39 | 165.00 | 38 | 20.52 | −56.25 | 58 | 25.65 | −45.00 |
| 19 | 15.39 | −165.00 | 39 | 20.52 | −33.75 | 59 | 25.65 | −27.00 |
| 20 | 15.39 | −135.00 | 40 | 20.52 | −11.25 | 60 | 25.65 | −9.00 |

ID ($i$), the identity of the POL unit referring to the figures of the main text; $\theta^i$, the zenith distance of the $i^{\text{th}}$ unit; $\phi^i$, the azimuth of the $i^{\text{th}}$ unit.
