## Supplementary material for "From skylight input to behavioural output: a computational model of the insect polarised light compass": S3 Appendix

**S1 Appendix.** **The objective function** used to verify the performance of the sensor combines a detailed simulation of the skylight, the POL-unit processing, the CX compass model that predicts the sun position, and the decoder of the output signal.

**Sky simulation.** To estimate the values of the photo-receptors we need to extract the intensity, degree and angle of polarisation of the sky for the pointing directions of all the units. For that reason we integrate a simulation of the sky in our objective function, which computes these features given the sun position.

For the simulation of the sky, we follow [1], who uses the [2] sky-dome relative luminance model, which is found to be more accurate than the widely cited CIE model [3]:

$$\mathcal{F}(z,\chi) = (1 + Ae^{B/\sin(z)})(1 + Ce^{D\chi} + E\cos^2(\chi))$$

where $A$, $B$, $C$, $D$ and $E$ are constant parameters that describe the sky features and rely only on the *turbidity* of the atmosphere, $T_L$. More specifically, $A$ controls the darkening or brightening of the horizon, $B$ the luminance gradient near the horizon, $C$ the relative intensity of the circumsolar region, $D$ the width of the circumsolar region, and $E$ the relative backscattered light. The parameters of the above function $z$ and $\chi$ are the angular distance of the sky element from the zenith point and the sun position respectively. For every element, $j$, in the sky-dome with elevation, $\theta_t^j$, and azimuth, $\phi_t^j$, we can get its absolute luminance, $L_t^j$ $(kcd/m^2)$, using:

$$L_t^j = L_z \frac{\mathcal{F}(\theta_t^j, \gamma_t^j)}{\mathcal{F}(\theta_s, 0^\circ)} \tag{1}$$

where $\gamma_t^j$ is the angular distance between the sky element $j$, tilted according to the tilting orientation $t$, and the sun position $s$, and is given by the equation:

$$\gamma_t^j = \cos^{-1}[\sin\left(\theta_t^j\right)\sin(\theta_s) + \cos\left(\theta_t^j\right)\cos(\theta_s)\cos\left(\phi_t^j - \phi_s\right)]$$

where $\theta_s$ and $\phi_s$ are the solar elevation and azimuth respectively. The zenith luminance, $L_z$ $(kcd/m^2)$, depends on the Linke's turbidity factor ($T_L$) and the distance of the sun from the zenith point, $\theta_s$, and is given by:

$$L_z = (4.8453 \cdot T_L - 4.9710) \cdot \tan\left[\left(\frac{4}{9} - \frac{T_L}{120}\right)(180^\circ - 2\theta_s)\right] - 0.2155 \cdot T_L + 2.4192$$

The polarisation pattern is highly influenced by the sky intensity. To accurately model the degree of polarisation we need to calculate this influence using:

$$I_t^j = \left[\frac{1}{\mathcal{F}(\theta_t^j, \gamma_t^j)} - \frac{1}{\mathcal{F}(\theta_s, 0^\circ)}\right] \frac{\mathcal{F}(\theta_s, 0^\circ) \cdot \mathcal{F}(||\theta_s - 90^\circ||, 90^\circ)}{\mathcal{F}(\theta_s, 0^\circ) - \mathcal{F}(||\theta_s - 90^\circ||, 90^\circ)}$$

Therefore, the degree of polarisation is:

$$P_t^j = \frac{1}{90^\circ} e^{-\frac{T_L - C_1}{C_2}} \frac{\sin^2(\gamma_t^j)}{1 + \cos^2(\gamma_t^j)} \left[\left(90^\circ - \theta_t^j\right)\sin\left(\theta_t^j\right) + \theta_t^j I_t^j\right] \tag{2}$$

where $C_1 = 0.6$ and $C_2 = 4.0$ are parameters that regulate the maximum degree of polarisation according to the turbidity. The angle of polarisation depend only on the positions of the sun and the observed element and is given by:

$$A_t^j = \tan^{-1}\left(\frac{x_t^j}{y_t^j}\right) \tag{3}$$

where $x_t^j$ and $y_t^j$ are the Cartesian coordinates of the polarisation vector, the angle of which on the $(x, y)$ plane is parallel to the angle of polarisation, and their values are given by:

$$x_t^j = \cos\left(\theta_t^j\right)[\sin\left(\phi_t^j\right) - 2\sin^2(\frac{\theta_s}{2})\sin(\phi_s)\cos\left(\phi_s - \phi_t^j\right)] - \sin\left(\theta_t^j\right)\sin(\theta_s)\sin(\phi_s)$$

$$y_t^j = \cos\left(\theta_t^j\right)\cos\left(\phi_t^j\right)[\cos(\theta_s)\cos^2(\phi_s) + \sin^2(\phi_s)] - \cos\left(\theta_t^j\right)\sin\left(\phi_t^j\right)\sin^2(\frac{\theta_s}{2})\sin(2\phi_s) - \sin\left(\theta_t^j\right)\sin(\theta_s)\cos(\phi_s)$$

**Signal encoding.** As described before, the output of the $j^{\text{th}}$ POL-OP unit, tilted by $t$, is given by:

$$\left(r_{\text{POL}}^j\right)_t = \frac{\left(r_\parallel^j\right)_t - \left(r_\perp^j\right)_t}{\left(r_\parallel^j\right)_t + \left(r_\perp^j\right)_t} \quad (4)$$

where $\left(r_\parallel^j\right)_t$ and $\left(r_\perp^j\right)_t$ are the responses of the polarisation-opponent neurons of the model in the position $j$ on the sensor, while the sensor is tilted by the $t$ pair of angles. Their values are given by:

$$\left(r_\parallel^j\right)_t = \sqrt{\left(s_\parallel^j\right)_t}, \qquad \left(s_\parallel^j\right)_t = L_t^j \cdot \left[\sin^2(A_t^j - \alpha_t^j) + \cos^2(A_t^j - \alpha_t^j)(1 - P_t^j)^2\right]$$

$$\left(r_\perp^j\right)_t = \sqrt{\left(s_\perp^j\right)_t}, \qquad \left(s_\perp^j\right)_t = L_t^j \cdot \left[\cos^2(A_t^j - \alpha_t^j) + \sin^2(A_t^j - \alpha_t^j)(1 - P_t^j)^2\right]$$

where $A_t^j$ is the angle of polarisation, $P_t^j$ is the degree of polarisation, and $\alpha_t^j = \phi_t^j - 90°$ is the corresponding orientation of the $j^{\text{th}}$ unit. For the computational model that transforms the POL-OP units responses into TCL responses, $r_{\text{TCL}}^k$, see methods.

**Signal decoding.** To decode the output of the computational model we use the Fast Fourier Transform (FFT). Therefore, the vector that points towards the sun is given by:

$$R = \sum_{k=1}^{n_{\text{TCL}}} r_{\text{TCL}}^k e^{-i \cdot 360°(k-1)/n_{\text{TCL}}}$$

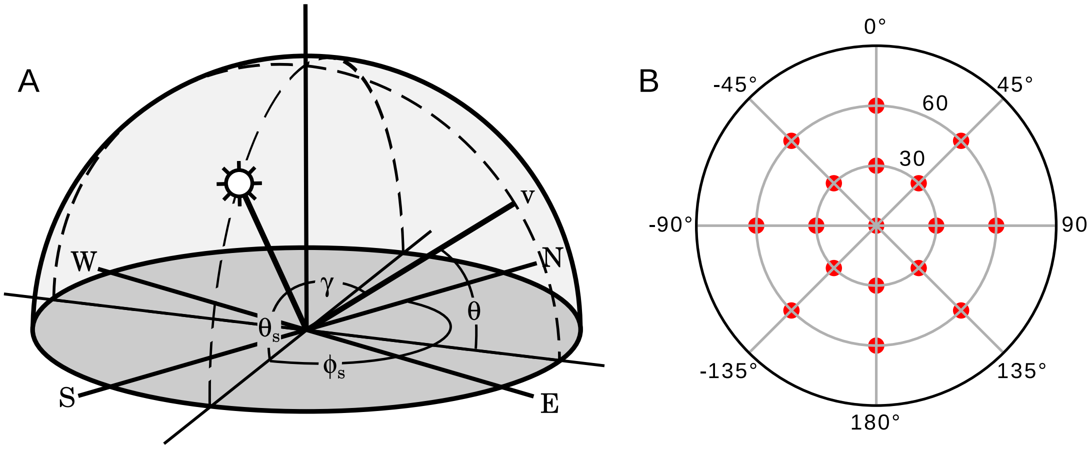


**Figure 1.** Visual representation of the parameters of the objective function. (A) Parameters extracted from the sun and element position on the sky-dome. (B) Tiling angles; maximum tilting angle: 60°; total number of tilting directions: 17.

where $R \in \mathbb{C}$. The angle of this complex number gives the solar azimuth, $\phi'_s$, while the magnitude imply the confidence of this prediction, $\tau_s = \frac{1}{\sigma_s}$:

$$\begin{aligned}\phi'_s &= \Phi(\theta_t^1, ..., \theta_t^n, \phi_t^1, ..., \phi_t^n, \alpha_t^1, ..., \alpha_t^n, \mathbf{W}, \theta_s, \phi_s) = 360° - \tan^{-1}\left[\frac{\mathrm{Im}(R)}{\mathrm{Re}(R)}\right] \\ \tau_s &= \Theta(\theta_t^1, ..., \theta_t^n, \phi_t^1, ..., \phi_t^n, \alpha_t^1, ..., \alpha_t^n, \mathbf{W}, \theta_s, \phi_s) = ||R||\end{aligned} \tag{5}$$

**Objective function.** We calculate the performance of a specific design and model using the angular error between the original and the predicted solar azimuth. In order to make the objective function invariant to the sun position, we compute the error over 500 sun positions homogeneously distributed in the sky-dome, i.e. using the Fibonacci spherical distribution, and 17 different tilting orientations. The total cost will then be the average value among these errors weighted by the confidence of the prediction. Hence,

$$J(\theta^1, ..., \theta^n, \phi^1, ..., \phi^n, \alpha^1, ..., \alpha^n, \mathbf{W}) = \frac{1}{8500}\sum_{t=1}^{17}\sum_{s=1}^{500}||[(\phi_s - \phi'_s + 180°) \mod 360°] - 180°||$$

gives an estimation of the angular error of the given design and computational model.
