## Supplementary material for "From skylight input to behavioural output: a computational model of the insect polarised light compass": S4 Appendix

**S2 Appendix. Integrating for infinite SOL-neurons.** The computational model can also be described as a linear combination of the outputs of the POL-OP units. We can merge the SOL and TCL layers, in one by integrating for infinite SOL-neurons. Therefore, the output of each TCL neuron is given by:

$$r_{\mathrm{TCL}}^{k} = \sum_{j=1}^{n_{\mathrm{POL}}} W^{j,k} \cdot g^{j}(\theta_t, \phi_t) \cdot r_{\mathrm{POL}}^{j}$$

where $W^{j,k}$ is the weight of the synapse connecting the input, $j$, and the output, $k$. The weights of the model are given by the equation below:

$$W^{j,k} = -\frac{n_{\mathrm{TCL}}}{2 \cdot n_{\mathrm{POL}}} \cdot g^{j}(\theta_t, \phi_t) \cdot \sin\left[\left(\phi_{\mathrm{TCL}}^{\mathrm{pref}}\right)^{k} - \alpha^{j}\right]$$

where $\left(\phi_{\mathrm{TCL}}^{\mathrm{pref}}\right)^{k}$ is the preference angle of the $k^{\mathrm{th}}$ TCL-neuron. This equation is the result of integrating the two processing steps (SOL and TCL) to one by assuming infinite SOL-neurons. The processing would look like this:

$$r_{\mathrm{TCL}}^{k} = \sum_{\xi=1}^{n_{\mathrm{SOL}}} W_{\mathrm{TCL}}^{\xi,k} \sum_{j=1}^{n_{\mathrm{POL}}} W_{\mathrm{SOL}}^{j,\xi} \cdot g^{j}(\theta_t, \phi_t) \cdot r^{j}$$

where

$$\begin{aligned}
g^{j}(\theta_t, \phi_t) &= \exp\left\{ -\frac{1}{2}\left[\frac{\sin\left(\theta_g - \theta^{j}\right)\cos(\theta_t) + \cos\left(\theta_g - \theta^{j}\right)\sin(\theta_t)\cos\left(\phi_t - \phi^{j}\right)}{\sigma_g}\right]^2\right\} \\
W_{\mathrm{SOL}}^{j,\xi} &= \frac{n_{\mathrm{SOL}}}{n_{\mathrm{POL}}} \cdot \sin\left[\alpha^{j} - \left(\phi_{\mathrm{SOL}}^{\mathrm{pref}}\right)^{\xi}\right] \\
W_{\mathrm{TCL}}^{\xi,k} &= \frac{n_{\mathrm{TCL}}}{2 \cdot n_{\mathrm{SOL}}} \cos\left[\left(\phi_{\mathrm{TCL}}^{\mathrm{pref}}\right)^{k} - \left(\phi_{\mathrm{SOL}}^{\mathrm{pref}}\right)^{\xi}\right]
\end{aligned}$$

Integrating for $\delta = \left(\phi_{\mathrm{SOL}}^{\mathrm{pref}}\right)^{\xi}$ in the limit where $n_{\mathrm{SOL}} \to +\infty$ we have:

$$\begin{aligned}
W^{j,k} &= \lim_{n_{\mathrm{SOL}} \to +\infty} \sum_{\xi=1}^{n_{\mathrm{SOL}}} W_{\mathrm{TCL}}^{\xi,k} W_{\mathrm{SOL}}^{j,\xi} \\
&= \lim_{n_{\mathrm{SOL}} \to +\infty} \frac{n_{\mathrm{TCL}}}{2 \cdot n_{\mathrm{POL}}} \sum_{j=1}^{n_{\mathrm{CBL}}} \cos\left[\left(\phi_{\mathrm{TCL}}^{\mathrm{pref}}\right)^{k} - \delta\right] \sin\left(\alpha^{j} - \delta\right) \\
&= \frac{n_{\mathrm{TCL}}}{2 \cdot n_{\mathrm{POL}}} \cdot \frac{1}{180^{\circ}} \int_{0}^{360^{\circ}} \cos\left[\left(\phi_{\mathrm{TCL}}^{\mathrm{pref}}\right)^{k} - \delta\right] \sin\left(\alpha^{j} - \delta\right) d\delta \\
&= -\frac{n_{\mathrm{TCL}}}{2 \cdot n_{\mathrm{POL}}} \cdot \sin\left[\left(\phi_{\mathrm{TCL}}^{\mathrm{pref}}\right)^{k} - \alpha^{j}\right]
\end{aligned}$$

Using this model we get almost the same result as the two-layered one.
